# Chromosomal rearrangements and segmental deletions contribute to gene loss in squamates

**DOI:** 10.1101/2025.07.03.663111

**Authors:** Buddhabhushan Girish Salve, H Sonal, Nagarjun Vijay

## Abstract

**Background:** Genomic rearrangements, including segmental deletions, duplications, translocations, and inversions of DNA segments, can contribute to gene losses, thereby reshaping genome architecture and potentially resulting in functional consequences. In squamates, karyotypic evolution mainly involves chromosome number reduction through fusions and microchromosome to macrochromosome translocations, although fissions have also contributed to diversification in several lineages. Despite these dynamics, the evolutionary processes and underlying genetic mechanisms driving chromosomal rearrangements and associated gene losses in squamates remain poorly understood.

**Results:** In this study, we analysed chromosome/scaffold-level assemblies of 261 squamates, corroborated by short-read, long-read, and transcriptomic data. We found multiple lines of evidence for the putative loss of 53 genes in the squamate lineage. Synteny and phylogenetic analysis revealed that, among the 53 unretrieved orthologs, 14 are lost in squamates with no retained paralog, 15 show ortholog loss with retained paralogs, and 24 remain as unretrieved orthologs. Furthermore, we find that many of the genes lost from squamates are organised in syntenic clusters and are involved in essential immune functions—raising important questions about the role of paralogs in compensating for the function of lost genes, strengthening the “less is more” hypothesis in the squamate lineage.

**Conclusions:** Together, our comparative genomic analyses highlight that the loss of crucial genes in squamate lineages has occurred primarily through inter- and intrachromosomal rearrangements, including segmental deletions. These findings offer insights into the evolutionary loss of genes involved in macrophage differentiation and inform the development of novel pharmaceutical approaches for modulating immune responses.

## Background

Genomic rearrangements, including deletions, inversions, translocations, and duplications, play a major role in shaping genome architecture and can potentially contribute to gene loss [1–3]. Such structural changes often disrupt gene integrity by breaking coding regions, altering regulatory landscapes, or relocating genes into heterochromatic or transcriptionally silent regions [4, 5]. For example, large-scale deletions can physically remove entire genes, whereas rearrangements such as translocations or inversions may separate essential gene components or place them under incompatible regulatory control, rendering them non-functional [3, 6, 7]. Moreover, rearrangements mediated by repetitive elements can create unstable genomic regions prone to further structural variation, thereby accelerating gene erosion [8–12]. These processes are particularly evident in rapidly evolving lineages, such as birds and mammals, where gene loss contributes to lineage-specific adaptation and diversification [13–16].

The recent dramatic increase in the number of high-quality genome assemblies of diverse organisms has facilitated studies on lineage-specific gene losses [17–20]. The NCBI RefSeq annotation of orthologues for multiple vertebrate species and the availability of short- and long-read sequencing data allow the confirmation of gene losses [21, 22]. However, most studies linking gene loss and genomic rearrangements in vertebrate evolution have focused on birds [3, 23], mammals [11] and fishes [24], and the association of such chromosomal rearrangement and gene loss have not been systematically performed in squamates.

Squamate reptiles are a diverse monophyletic group of vertebrates classified into lizard and snake families, which diverged from tuatara approximately 250 million years ago (MYA) [25, 26]. With more than 11,000 described species, they are more species-rich than birds [27]. Squamates have emerged as excellent systems for studying genomic rearrangement [28], karyotype [29], genome size evolution [30], and unique biology such as tissue regeneration and limb degeneration [31, 32]. Yet, the dearth of genome assemblies has been a bottleneck for large-scale comparative genomic studies [33]. The recent availability of high-quality genomes across the squamate tree of life has opened new avenues to look at lineage-specific genome dynamics, such as sex chromosome evolution [34], changes in the repeat landscape [35], and GC content heterogeneity [36]. Notably, the genome size of squamates is heterogenous, with a trend toward genome size reduction [37]. Similar reductions in birds and bats have been associated with the loss of gene families and entire syntenic blocks, suggesting that the contraction of the gene repertoire may accompany genome size reduction [37, 38]. Although gene and whole-genome duplications drive evolutionary innovation, emerging evidence now reinforces the “less-is-more” hypothesis, highlighting gene loss as an equally potent force in shaping genomic and phenotypic diversity [39–41]. The growing availability of chromosome-scale assemblies from initiatives such as the Vertebrate Genomes Project (VGP) is now enabling the systematic identification of such gene loss events, providing unprecedented opportunities to explore how gene reduction contributes to lineage-specific adaptation and evolutionary novelty [42–44]. In this context, investigating patterns of gene loss across the squamate radiation can therefore yield critical insights into how genomic reduction and restructuring contribute to phenotypic innovation, evolutionary diversification, and an alternative to phyletic gradualism [45].

To address this gap, we focused on gene loss in squamates and the role of genomic rearrangements. Our preliminary genome-wide screening of vertebrates revealed that 53 key genes with conserved functions are not annotated in NCBI RefSeq in squamates. Among these genes, previous studies have reported the loss of *PLAAT1*, *SLC24A1,* and *IL34* in squamates and associated them with adaptation to low-light environments [46], changes in the visual system [47], and myeloid cell-type evolution [48], respectively. These studies suggest the evolutionary loss of these genes based on a search of the genome assemblies and failure to recover any remnants. Here, we leverage high-quality genomes of squamates coupled with a comprehensive search of genomic short- and long-read and transcriptomic data, assembly validation, syntenic and phylogenetic analysis to rule out inadequacies in the genome assembly posing as gene loss [42, 49, 50]. We began our search by looking for genes annotated in the five outgroup species (i.e., Western clawed frog (*Xenopus tropicalis*), American alligator (*Alligator mississippiensis*), chicken (*Gallus gallus*), green sea turtle (*Chelonia mydas*), and human (*Homo sapiens*)) and not annotated in all annotated squamate genomes. Furthermore, we looked for direct (search of genomes, raw read datasets, statistical and phylogenetic signal of gene loss) and indirect evidence (co-evolution of the receptor) supporting the loss of these genes. We found that gene loss in squamates occurred in conserved syntenic clusters, similar to the pattern observed in birds [38]. The putative gene loss we identified needs to be evaluated with better-quality genome assemblies, and its functional impact should be assessed through comparative functional assays.

## Results

First, as a pilot analysis to identify unretrieved orthologs specific to the entire squamate lineage, we focused on genes consistently identified as not annotated in NCBI RefSeq across all 30 annotated squamate genomes, while being present in outgroup species (Additional file 1: Table S1-S2). To ensure robust inference, we used high-quality genome assemblies from multiple vertebrate species, including 261 squamates. Of these, 77 assemblies meet the quality standards recommended by global initiatives such as the Vertebrate Genome Project and the Earth BioGenome Project, with contig N50 values exceeding 1 Mb and scaffold N50 values exceeding 10 Mb (Additional file 1: Table S3). The 53 unretrieved orthologs were systematically classified into distinct categories based on multiple lines of genomic and syntenic evidence. These included: (i) assembly verification using long-read data through the Klumpy tool, (ii) local synteny assessment, (iii) evidence for segmental deletion or chromosomal rearrangement, (iv) ortholog sequence properties such as GC-content distribution and sequence identity with human orthologs, (v) local assembly using HybPiper followed by phylogenetic analysis, (vi) pairwise genome alignment, (vii) paralog annotation, (viii) transcriptional activity at the syntenic locus and cross-species expression mapping in central bearded dragon and chicken, (ix) BLASTn and tBLASTx searches in the near T2T phased genome assembly of central bearded dragon, and (x) BLASTn search against 261 squamate genomes. Based on the integration of syntenic and phylogenetic evidence, unretrieved orthologs were further subdivided into three categories: (i) ortholog loss with no retained paralog, supported by evidence for segmental deletion and/or syntenic and phylogenetic analysis and GC-content < 65% and sequence identity > 55% with no sequence paralog found, (ii) putative ortholog loss with retained paralog, syntenic and phylogenetic analysis found paralog along with loss of orthologs, and (iii) unclear cases, where high sequence divergence for highly diverged ortholog or translocated ortholog with high sequence divergence or germline-restricted genes or hard to sequence genomic regions containing unretrieved orthologs present in dark matter or non-B-form DNA structures or proximity to telomeric and centromeric regions, prevented confident classification (Additional file 1: Table S1 and Additional file 2: Gene-wise evidences).

### Unretrieved orthologs occur in conserved syntenic blocks and chromosomal rearrangements

We examined the genomic locations of the unretrieved orthologs in the chicken genome and found that they are distributed across both macro- and microchromosomes (**Fig. 1**). This contrasts with previous studies on gene loss in birds, where unretrieved orthologs were predominantly located on microchromosomes [38, 42]. We found clusters of unretrieved orthologs on chromosomes 1 (*LCP1*, *RUBCNL*, *SIAH3*, *LRCH1*, *ARL11*, *DLEU7*, *STOML3*, *INTS6* and *FREM2*) and 4 (*TBC1D14*, *MAN2B2*, *GPR78*, *GRK4*, *TNIP2*, *KDM3A*) of chicken. We also noted 7 clusters of two or more genes within ∼ 2 Mb of each other in the chicken genome. The cross-species comparison of syntenic loci revealed that these regions are prone to both intra- and inter-chromosomal rearrangements in squamate species such as the central bearded dragon (*Pogona vitticeps*), brown anole (*Anolis sagrei*), and terrestrial garter snake (*Thamnophis elegans*) (**Fig. 1**; Additional file 1: Table S4). In several cases, these loci are also mapped to telomeric regions. In contrast, comparable rearrangements were not observed in other reptiles, including the American alligator (*Alligator mississippiensis*) and the green sea turtle (*Chelonia mydas*), suggesting squamate-specific genomic rearrangements.

**Fig. 1.**
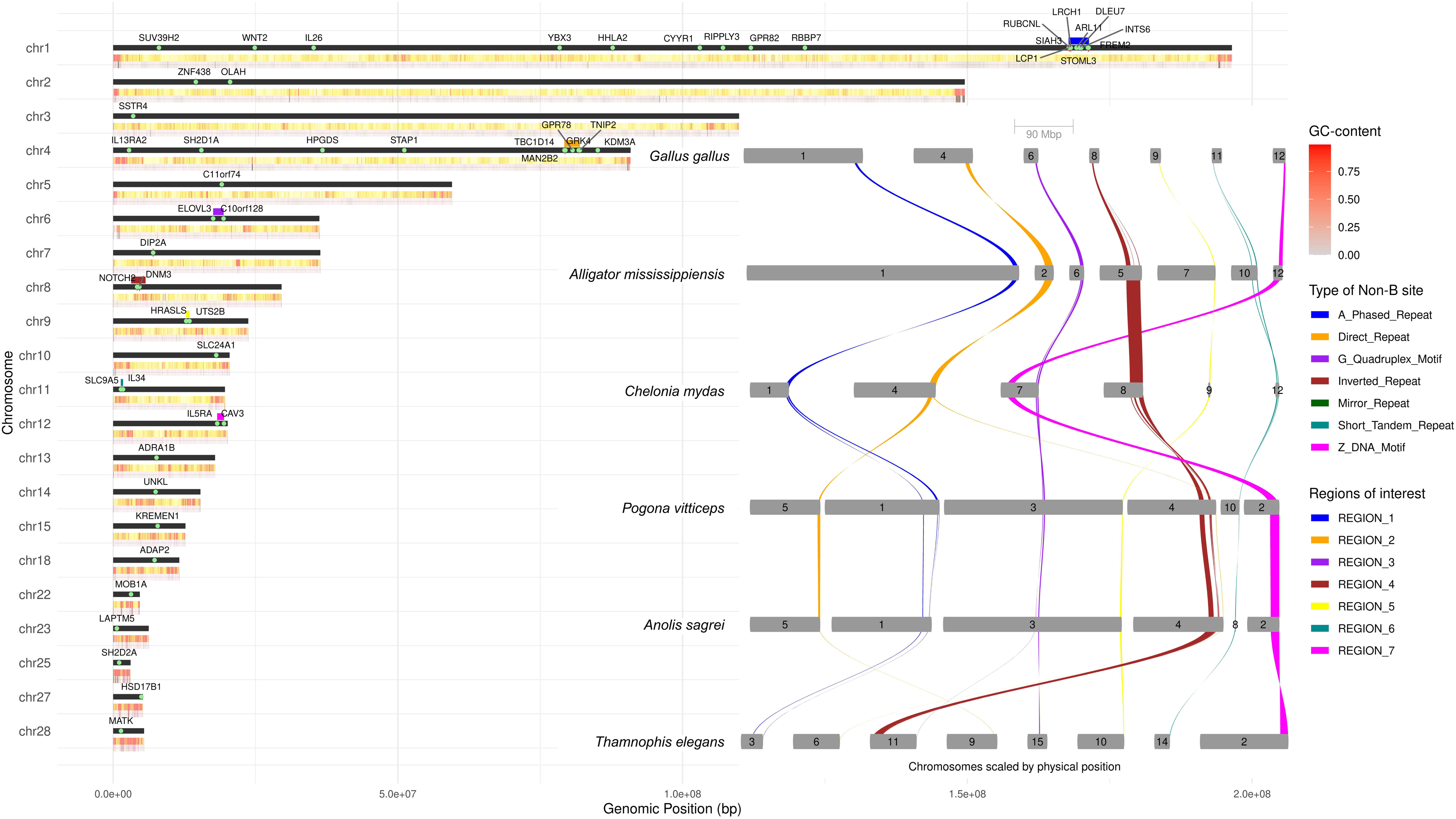
Unretrieved orthologs are localised to discrete chromosomal regions, occur within syntenic blocks, and are associated with chromosomal rearrangements. The left panel shows the genomic locations of 53 unretrieved orthologs mapped across 21 chromosomes of the Chicken karyotype (black horizontal bars scaled by genomic position on the x-axis, and chromosome numbers are shown on the left side). Unretrieved orthologs are indicated by light green dots labelled with gene names. GC content, calculated in 100 bp windows, is represented by a gradient from light grey (low GC) to red (high GC). Below the GC track, the presence of predicted non-B DNA motifs is indicated by colour-coded bars. The right panel depicts chromosomal rearrangements across representative sauropsid species (*Gallus gallus*, *Alligator mississippiensis*, *Chelonia mydas*, *Pogona vitticeps*, *Anolis sagrei*, and *Thamnophis elegans).* The chromosome numbers are mentioned above each grey-coloured rectangle, and coloured ribbons (as defined in the left panel) represent conserved synteny blocks connecting orthologous regions, with seven regions of interest (ROI_1 to ROI_7) highlighted. These regions are implicated in the loss of two or more genes. The chicken orthologues of the human genes *IFTAP*, *PLAAT1*, and *TMEM273* correspond to *C11orf74*, *HRASLS*, and *C10orf128*, respectively.

### Integrated multi-evidence stratified framework for putative gene loss validation

Based on the syntenic analysis along with local genome assembly validation, the 53 unretrieved orthologs could be classified as putatively lost due to segmental deletions (n=36) and intra/inter chromosomal rearrangements (n=17) (**Fig. 2A**). In the case of segmental deletions, the evidence for the loss of the gene consists of squamate-specific phylogenetic signal of reduction in intergenic distance, genome blast coverage, and phylogenetic assessment of sequences assembled by HybPiper or retrieved by blast search. While in the case of chromosomal rearrangements, the evidence relied largely on syntenic and phylogenetic analysis. An evidence score for the loss of each gene based on these integrated multiple layers of evidence was used to rank the unretrieved orthologs.

**Fig. 2.**
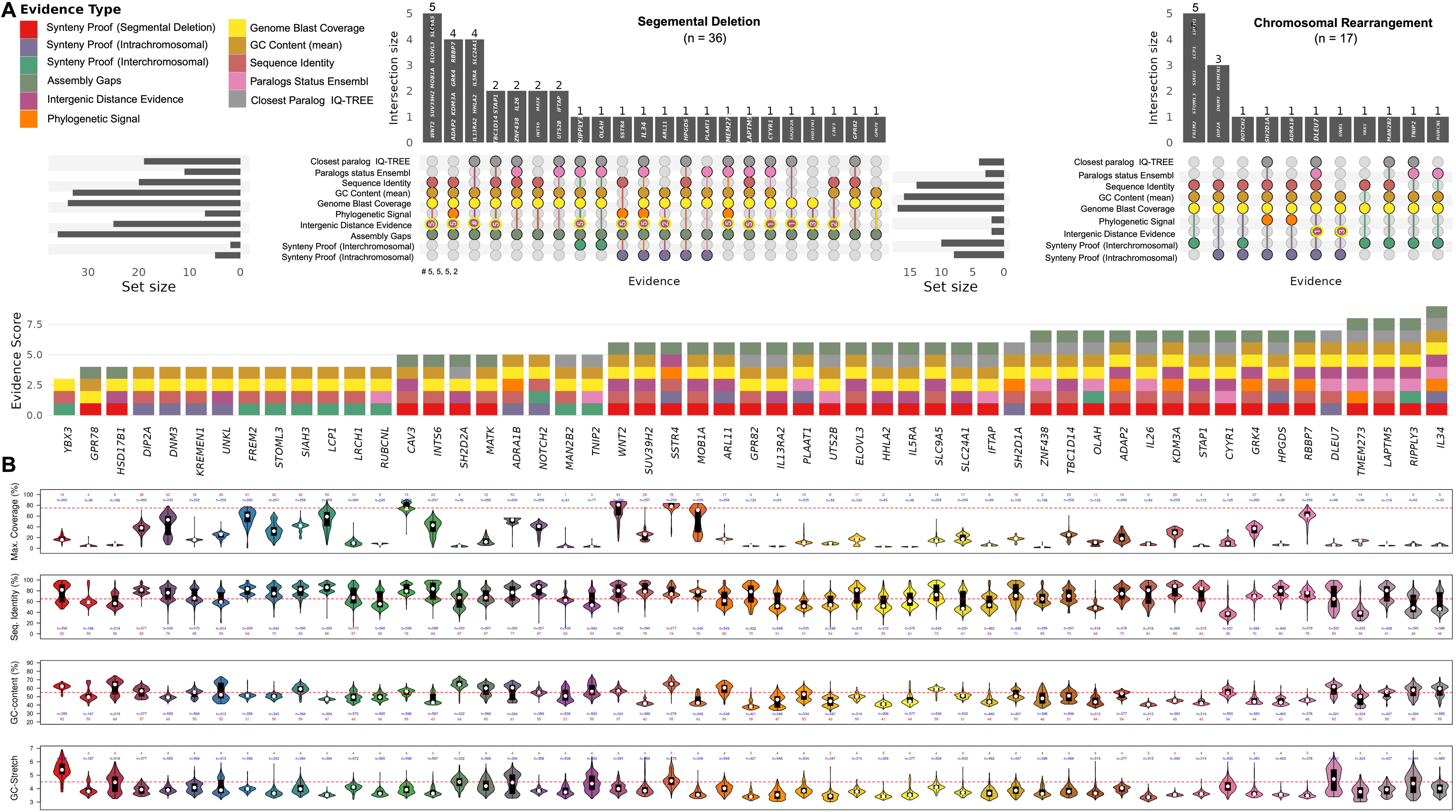
Multiple lines of evidence support gene loss in the Squamate lineage. **A.** Evidence types used to validate candidate gene losses/unretrieved orthologs. The left panels summarise evidence of segmental deletions (n = 36 genes) and chromosomal rearrangements (n = 17 genes). Upset plots display intersections of evidence categories, including synteny, genome blast coverage, GC content, sequence identity, paralog status (Ensembl), closest paralog placement (Hybpiper and IQ-TREE-based phylogenetic analysis), phylogenetic signal for segmental deletion, intergenic distance-based evidence of segmental deletion, and assembly gaps found using klumpy. Horizontal bar plots show total set size for each evidence type, while vertical bars represent intersection sizes and gene names are mentioned on bars (the font size varies with increase in evidence). The stacked bar plots below indicate cumulative evidence scores for each gene, with contributions color-coded by evidence type. **B.** Quantitative evaluation of evidence metrics across 53 candidate gene losses/unretrieved orthologs. Violin plots display distributions of (i) maximum genome coverage obtained using BLASTn search for each squamate genome, (ii) sequence identity (%), (iii) GC content (%), and (iv) GC stretches. Each gene is represented by its own distribution profile, with dashed lines marking thresholds used to support gene classification as candidate gene losses or unretrieved orthologs.

The BLASTn search of 261 squamate genomes revealed that most unretrieved orthologs are absent from squamates. Of the 53 genes, 21 have query coverage (search on squamate genomes using human orthologues as a query) of ≤ 25%, 23 genes between 25 - 74%, and 9 genes have coverage of ≥ 75% (*SUV39H2*, *FREM2*, *DNM3*, *RBBP7*, *MOB1A*, *SSTR4*, *WNT2*, *LCP1* and *CAV3*; **Fig. 2B**; Additional file 1: Table S5). Our bioinformatic analysis (using BioMart human paralog annotation) revealed two or more paralogs for each gene having query coverage >75% (**Fig. 3**; Additional file 1: Table S6). Furthermore, we assembled gene sequences using raw whole-genome sequencing datasets. The phylogenetic analysis of the retrieved sequences revealed that they cluster with closely related paralogues rather than the expected orthologues (Additional file 1: Table S7 and Additional file 2: Gene-wise evidences).

**Fig. 3.**
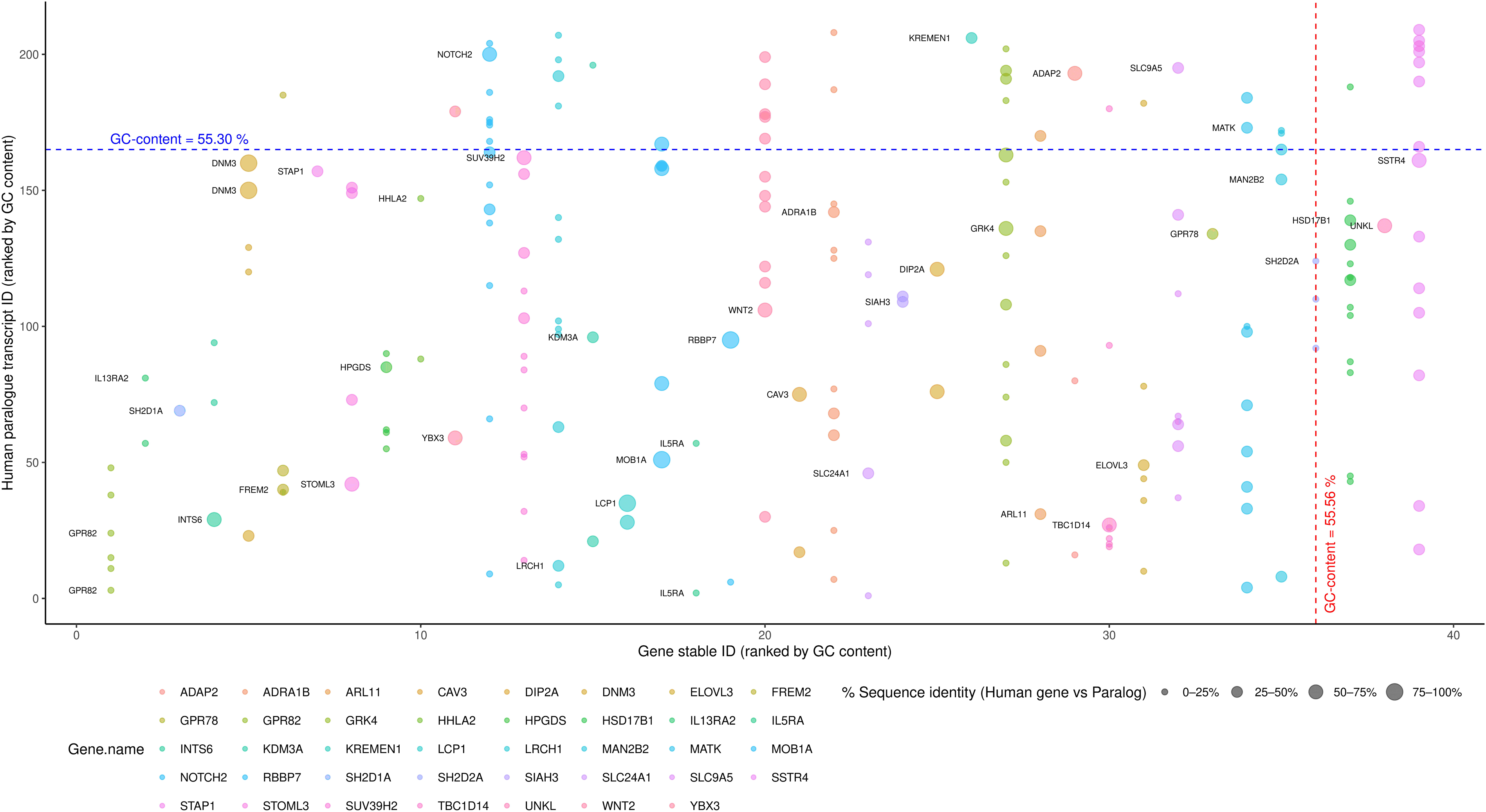
Unretrieved orthologs often belong to multi-gene families of paralogues. Human genes corresponding to squamate-unretrieved orthologs are plotted based on their GC-content (x-axis) and paralogs (y-axis) as annotated in Ensembl. Each point represents a paralogue, with the size of the light-coloured filled circle indicating the percentage sequence identity between the unretrieved orthologs and their paralog. Dashed lines indicate an approximate GC content of 55% for both axes (vertical red for the unretrieved orthologs and horizontal blue for the paralogues). This representation highlights that genes lost in squamates often belong to multi-gene families with moderate GC content.

It has been previously reported that unretrieved orthologs or “missing” or “hidden” genes in genome assemblies are often associated with high GC content and high sequence divergence [42, 49, 51, 52]. However, in our analysis, the mean GC content of the unretrieved ortholog (calculated across vertebrate orthologues) ranged from 39.56% to 64.72% (**Fig. 2B**). Notably, nineteen genes—*CAV3* (55.21%), *NOTCH2* (55.30%), *CYYR1* (55.43%), *DIP2A* (55.47%), *UNKL* (55.80%), *RIPPLY3* (56.06%), *ADRA1B* (56.48%), *IL34* (56.76%), *WNT2* (56.90%), *TNIP2* (57.83%), *SLC9A5* (58.12%), *MATK* (58.23%), *SIAH3* (58.52%), *DLEU7* (58.67%), *ARL11* (58.92%), *HSD17B1* (61.35%), *YBX3* (61.65%), *SH2D2A* (64.47%), and *SSTR4* (64.72%)—had GC content exceeding 55% (Additional file 1: Table S1).

Immune genes often evolve under strong selective pressure due to the ongoing evolutionary arms race between the host immune system and invading pathogens. This rapid evolution results in low amino acid conservation among orthologous proteins across species, limiting the effectiveness of sequence similarity-based tools such as BLASTn for detecting these genes [52–55]. Therefore, some of these genes may not be truly lost but may be present as highly diverged orthologs. We observed that 24 genes including *ARL11*, *GPR78, HHLA2, HPGDS, HSD17B1, IFTAP, IL13RA2, IL26, MATK, MOB1A, OLAH, PLAAT1, RBBP7, SH2D2A, SLC24A1, TMEM273, UTS2B, ZNF438, DLEU7, DNM3, UNKL, MAN2B2,*

*RUBCNL, and TNIP2* share less than 65% sequence identity with their human orthologs (**Fig. 2B**; Additional file 1: Table S1), suggesting that moderate to high sequence divergence may contribute to their loss or lack of annotation in squamate genomes, therefore remained classified as unretrieved orthologs. Comparative genome alignment between human and tuatara reveals that several genes were intact in the ancestral lineage with conserved gene synteny, highlighting their loss as specific to squamates (Additional File 1: Table S8).

Based on syntenic analysis, 36 genes were found to have a conserved gene order similar to that of the outgroup species, while eight genes showed evidence of intrachromosomal rearrangements and nine genes indicated interchromosomal rearrangements (**Fig. 2A**; Additional file 1: Table S1). Most of the 53 unretrieved orthologs lacked evidence of transcription, with some genes having spurious mapping (Additional File 3: Cross species RNA-seq mapping). We found evidence for segmental deletion in the case of 19 out of 36 genes, namely, *ADAP2*, *ELOVL3*, *GRK4*, *HHLA2*, *HSD17B1*, *IL34*, *IL5RA*, *KDM3A*, *LAPTM5*, *MOB1A*, *RBBP7*, *RIPPLY3*, *SLC24A1*, *SLC9A5, SSTR4, STAP1, SUV39H2, TBC1D14, TMEM273* and *WNT2* (Additional file 1: Table S1).

Paralogs were not found for 14 of the 53 unretrieved orthologs. Among these 14 genes, (i) *IL34*, *IL5RA*, and *RIPPLY3* exhibit high sequence divergence while retaining conserved synteny, with additional evidence of segmental deletion, (ii) A similar pattern (except for high sequence divergence) was observed for *GRK4*, *LAPTM5*, *SLC9A5*, *SSTR4*, *STAP1*, and *WNT2*, (iii) Despite weak evidence of segmental deletion, *CYYR1* (with frame-disrupting changes in *Anolis sagrei*, partial exon remnants in a genome assembly validated syntenic locus in *Pogona vitticeps* and lack of transcription in *Pogona vitticeps* and *Rhineura floridana*) and *GPR82* (with no blast hits even to paralogs despite low sequence divergence among orthologs) have conserved synteny (Additional file 4: Figure S1), (iv) The phylogenetic analysis suggests the lack of paralogs among the chromosomal rearrangement genes, *DIP2A*, *NOTCH2*, and *SH2D1A*. Collectively, these 14 genes were classified as cases of ortholog loss with no retained paralog (Additional file 1: Table S1, and Additional file 2: Gene-wise evidences).

Multi-gene families are often prone to inactivation or deletion, consistent with the birth-and-death model of gene family evolution [13, 24, 56, 57]. We found that 39 of the 53 unretrieved orthologs belong to multi-gene families and have at least one paralogue (**Fig. 2** and **Fig. 3**; Additional file 1: Table S1, S6 and S7). Among these 39 genes, (i) the evidence for conserved synteny and segmental deletion with limited sequence divergence was found for *ADAP2*, *CAV3*, *EVOVL3*, *INTS6*, *KDM3A*, *SUV39H2*, and *TBC1D14*, (ii) chromosomal rearrangement was identified in the case of *ADRA1B*, *KREMEN1*, *FREM2*, *LCP1*, *LRCH1*, *SIAH3*, *STOML3*, and *YBX3* with limited sequence divergence. Collectively, these 15 genes were classified as cases of ortholog loss with retained paralog (Additional file 1: Table S1, and Additional file 2: Gene-wise evidences).

Of the remaining 24 of the 39 genes with paralogs, *ARL11*, *GPR78*, *HHLA2*, *HSD17B1*, *IFTAP*, *IL13RA2*, *IL26, OLAH*, *PLAAT1*, *SH2D2A*, *SLC24A1*, *TMEM273*, *UTS2B*, *DLEU7*, *UNKL*, *FREM2*, *LCP1*, *LRCH1*, *MAN2B2*, *RUBCNL*, *SIAH3*, *STOML3*, and *TNIP2* had high sequence divergence and remain as unretrieved orthologs. In summary, synteny and phylogenetic analyses suggest that, among the 53 unretrieved orthologs, 14 genes are lost in squamates with no retained paralog, 15 show ortholog loss with retained paralogs, and 24 remain as unretrieved orthologs (Additional file 1: Table S1, and Additional file 2: Gene-wise evidences).

Furthermore, pair-wise genome alignments (at these 53 unretrieved orthologs loci) between chicken (*Gallus gallus*) and 34 other vertebrate species revealed that the squamate (∼280 MYA) genomes consistently having lower aligned exonic regions compared to tuatara (*Sphenodon punctatus*; ∼280 MYA), Testudines (∼261 MYA), and Crocodilians (∼245 MYA), despite these groups having comparable evolutionary divergence from chicken (**Fig. 4A**; Additional file 1: Table S9 and Additional file 4: Figure S2; pair-wise divergence time obtained from Timetree website [58]). The flanking genes to focal genes show significantly higher levels of alignments at exonic regions (**Fig. 4B**; Additional file 1: Table S9, and Additional file 2: Gene-wise evidences). The ENSEMBL available and chains files (pair-wise genome-wide LASTZ alignment) created by this study do not show any marked difference (Wilcoxon, p-value = 0.16, **Fig. 4C**). We also found that the aligned regions at exonic positions of focal genes are significantly lower in squamates compared to Birds, Coelacanthiformes, Testudines, Crocodylia, and Sphenodontia (**Fig. 4D**, Additional file 1: Table S9). The occurrence of unaligned regions in non-squamate species for some genes could be due to assembly gaps and/or high sequence divergence. The aligned region could also be from the paralogous regions.

**Fig. 4.**
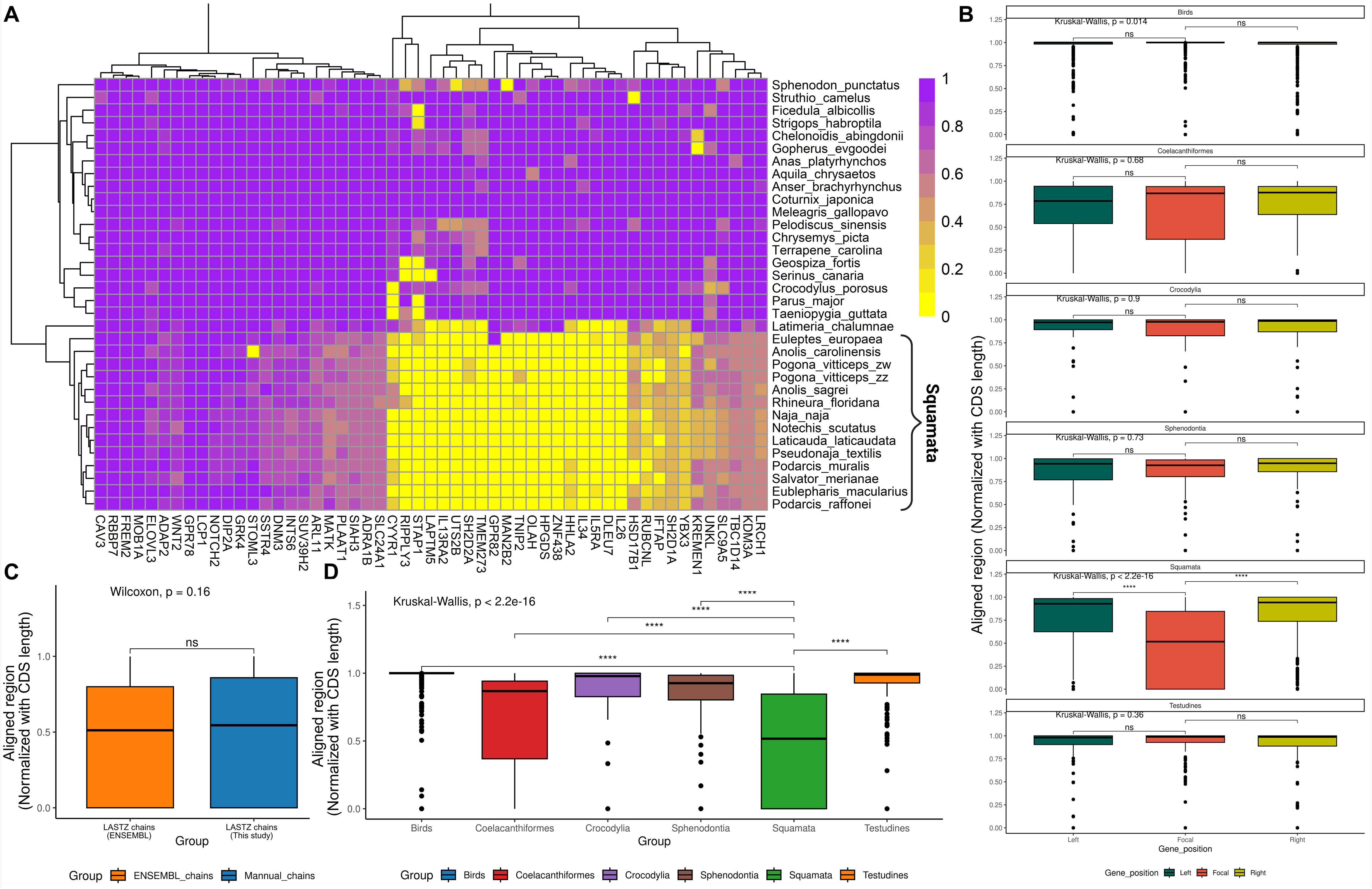
LASTZ alignment-based evidence for candidate gene loss in squamates. **A.** Heatmap of normalised aligned regions at the exonic region of Chicken (alignment length/CDS length) for 53 genes across 34 amniote species. Squamates show consistently reduced alignment at focal loci compared to other clades. In some cases, where alignment is detected, it may originate from paralogous sequences rather than true orthologs, while unaligned regions may reflect highly divergent orthologs. **B.** Boxplots of normalised aligned regions across clades for left, focal, and right gene positions. Significant reductions (wilcox.test; alternative = “greater”) are observed at focal loci in squamates, which is not the case for other orders. **C.** Comparison of Ensembl LASTZ chains (n = 7 species) and manually created LASTZ chains in this study (n = 6 species with two genomes of *Pogona vitticeps*, of squamate species BUSCO >97) shows no significant difference (Wilcoxon test, p = 0.16). **D.** Clade-level comparison highlights significantly lower alignment coverage in squamates relative to birds, crocodilians, testudines, and other lineages (wilcox.test; alternative = “greater”). Statistically significant pairwise differences are indicated by bars above the boxplots (*□p□≤□0.05, **□p□≤□0.01, ***□p□≤□0.001, ****□p□≤□0.0001, ns□=□non-significant). Boxplots are generated using ggplot [124].

### Comparative genomic evidence for *IL34* loss via segmental deletion in squamates

As reported previously [48], our comparative genomic analysis confirmed the absence of the *IL34* gene in squamates despite its widespread conservation across other vertebrates. Synteny analysis revealed that *COG4* and *SF3B3* typically flank *IL34* on the left and *MTSS2* and *VAC14* on the right (**Fig. 5A**; Additional file 1: Table S4). Notable exceptions are observed in species such as the zebrafish (*Danio rerio*), green sea turtle (*Chelonia mydas*), American alligator (*Alligator mississippiensis*), and tiger rattlesnake (*Crotalus tigris*), which have *IONP2*, *PARD6A*, *BBS2*, and *CNOT1,* respectively, on the right flank of the *IL34* locus. The lack of annotation for the *IL34* gene in squamates could potentially result from assembly artefacts. To rule out this possibility, we examined the syntenic locus of *IL34* in squamate species such as the common wall lizard (*Podarcis muralis*) and the Indian cobra (*Naja naja*) using PacBio and Oxford Nanopore long-read data. In both cases, overlapping reads span the *IL34* gene locus, indicating that the assemblies are correct at this locus (**Fig. 6**). BLASTn analysis of squamate genome assemblies and raw read datasets failed to recover any sequence corresponding to the *IL34* gene, further supporting its absence in this lineage and ruling out the possibility of its translocation to another genomic region (Additional file 1: Table S5 and S7). Next, we compared intergenic distances by examining squamate species for evidence of segmental deletions compared to non-squamate species (see Methods). Wilcoxon rank-sum tests (alternative = “less”) consistently yielded adjusted *p*-values < 0.001 for all combinations of gene distance ratios involving *SF3B3*–*MTSS2*, indicating significant contraction in this region in squamates (**Fig. 5B**; Additional file 1: Table S1; Additional file 4: Fig. S3). In contrast, other gene combinations did not significantly differ across all comparisons. Additionally, evidence for segmental deletion was supported by phylogenetic logistic regression analyses, with significant results from both logistic_MPLE (slope = 0.2; *p*-value = 0.0022) and logistic_IG10 (slope = 0.2; *p*-value = 0.0018) (**Fig. 5C**; Additional file 1: Table S1). Additionally, we found intrachromosomal rearrangement of the *IL34* locus in some of the Viperidae snake species (Additional file 4: Fig. S4).

**Fig. 5.**
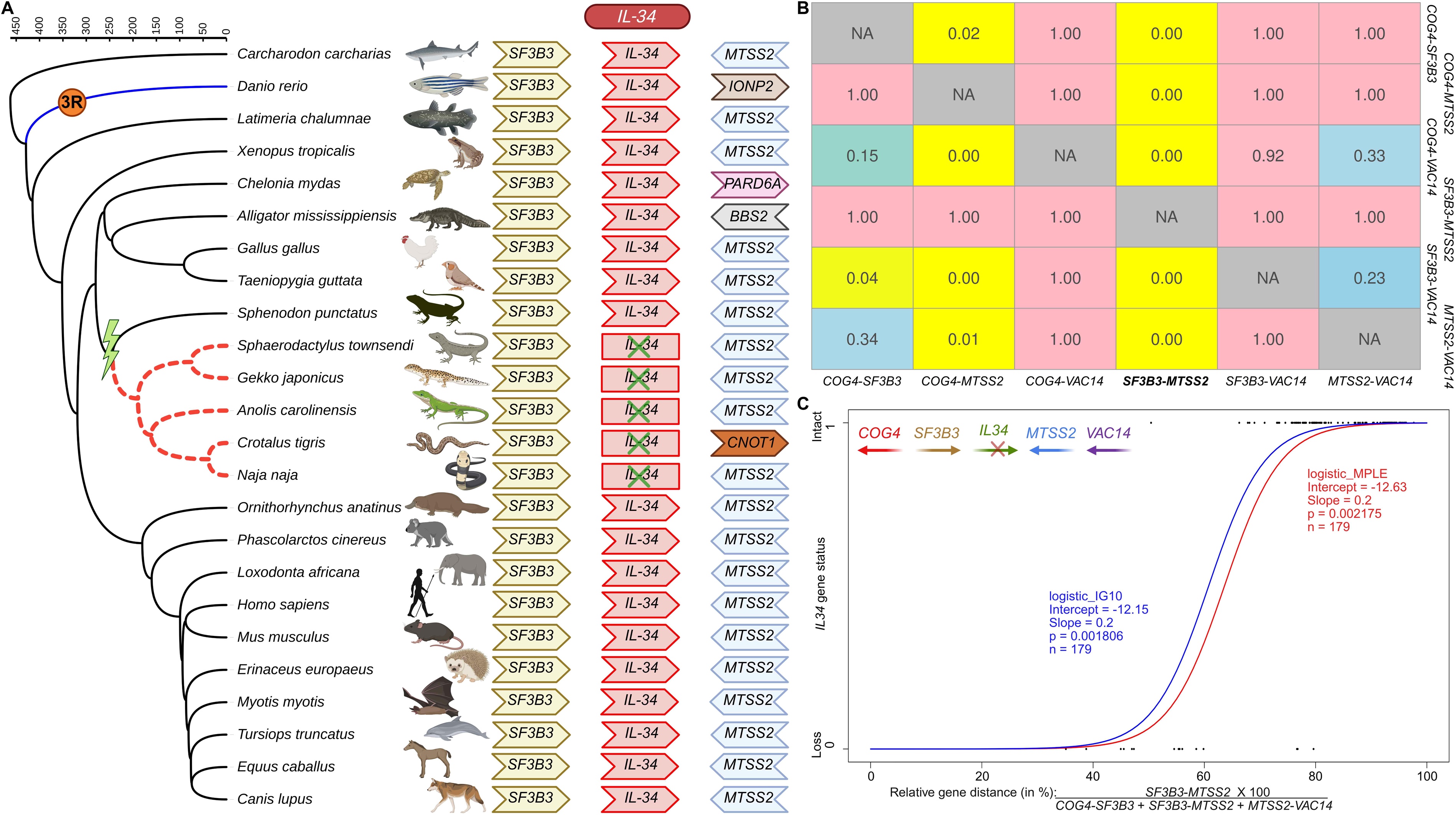
Evidence for *IL34* gene loss via segmental deletion. **A.** Micro-gene synteny of *IL34* in 24 vertebrate representative species. The phylogenetic branches of species with *IL34* lost are depicted with dashed red, while blue phylogenetic branches represent the 3^rd^ round of whole genome duplication. The green-coloured thunderbolt marks the gene loss event in squamates. Arrows represent genes, with their direction indicating gene orientation and gene names labelled inside each arrow. **B.** The heatmap shows the reduction in gene distance at the *IL34* locus (between *SF3B3*-*MTSS2*) in squamates. The values inside the box represent adjusted p-values (significant differences between squamates and non-squamates, calculated using pairwise Wilcox one-tailed test (alternative = “less”). **C.** Phylogenetic logistic regression analyses. The relative genomic distance between flanking genes (x-axis, in %) and the status of the *IL34* gene (y-axis) are correlated using two methods (logistic_IG10 and logistic_MPLE) to evaluate the segmental deletion of the *IL34* gene across vertebrates (n□=□179). The species tree was obtained from the Timetree website (https://timetree.org/) and annotated in iTOL (https://itol.embl.de/). Species images are from https://BioRender.com.

**Fig. 6.**
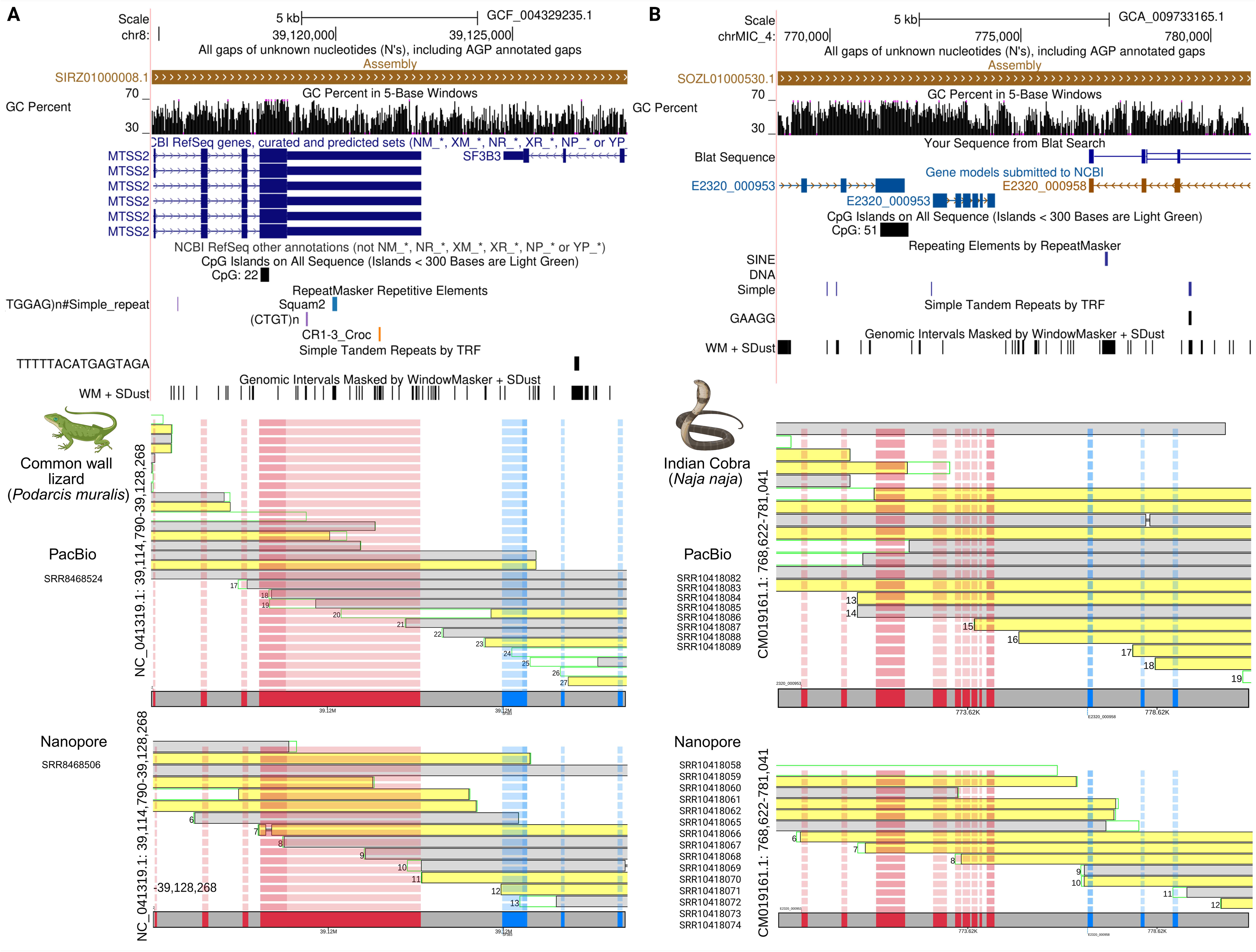
Validation of genome assembly at *IL34* gene syntenic locus in common wall lizard and Indian cobra genomes using long-read sequencing. UCSC Genome Browser snapshots of the *IL34* syntenic region are shown for the common wall lizard (*Podarcis muralis*; panel A, chr8:39,120,000–39,125,000) and the Indian cobra (*Naja naja*; panel B, chr4:770,000–780,000). The top portion of each panel displays GC content (black line), annotated syntenic genes (*MTSS2*/*SF3B3* in *P. muralis*, E2320_000953/958 in *N. naja*), and RepeatMasker annotations (including SINEs, LINEs, and low-complexity elements). The bottom portion of both panels shows long-reads from PacBio and Nanopore spanning the entire region. Grey and yellow segments indicate forward- and reverse-strand alignments, respectively. Transparent red and blue boxes mark exons of flanking syntenic genes. SRA (Short Read Archive) accession numbers for the datasets used to generate the BAM files are indicated next to the plots.

Our bioinformatic analysis, integrating synteny, long-read mapping-based genome assembly verification, BLASTn searches, transcriptome searches, intergenic distance comparisons, and phylogenetic regression, robustly confirms the loss of the *IL34* gene in squamates.

### Co-evolution of *CSF1R* with *IL34* gene loss in the squamate lineage

Our previous results from multifaceted approaches concurrently infer loss of the *IL34* gene from squamates. Next, we investigated the evolutionary consequences of *IL34* gene loss on its receptor —*CSF1R,* in squamates. Previous studies have suggested that *IL34* remains relatively unchanged, with detectable co-variation with *CSF1R*, whereas no such correlation is observed for *CSF1* [Garceau et al. [59], reviewed in [60]]. However, this hypothesis remains untested in lineages where *IL34* is lost. The availability of high-quality genome assemblies of squamates and *IL34* gene loss opens the opportunity to address this interesting evolutionary question─how two ligands evolve with a single receptor? Using selection analysis tools, we detected signatures of relaxed selection acting on *CSF1R* in several squamate species. We found squamate species such as Western terrestrial garter snake (*Thamnophis elegans*; *p*-values = 0.0003, k-value = 0.63), komodo dragon (*Varanus komodoensis*; *p*-value = 0.0053, k-value = 0.62), and central bearded dragon (*Pogona vitticeps*; *p*-value = 0.0047, k-value = 0.59) show signatures of relaxed selection by RELAX (with *p*-values <0.05 and k-value <0) and/or codeml (**Fig. 7**; Additional file 1: Table S10). Notably, signatures of relaxed selection were not pervasive across the screened vertebrate lineages, suggesting lineage-specific evolutionary changes. Several sites within the extracellular domain of *CSF1R* were identified as positively selected by both MEME and FEL analyses, with squamate species exhibiting a relatively higher number of positively selected sites (Additional file 1: Table S10). Together, the results of comparative sequence analyses revealed lineage-specific modifications in *CSF1R* in squamates, potentially reflecting compensatory evolution following *IL34* gene loss. These findings provide indirect but compelling evidence for *IL34* gene loss in squamates and support the hypothesis of receptor adaptation following the loss of one of its ligands.

**Fig. 7.**
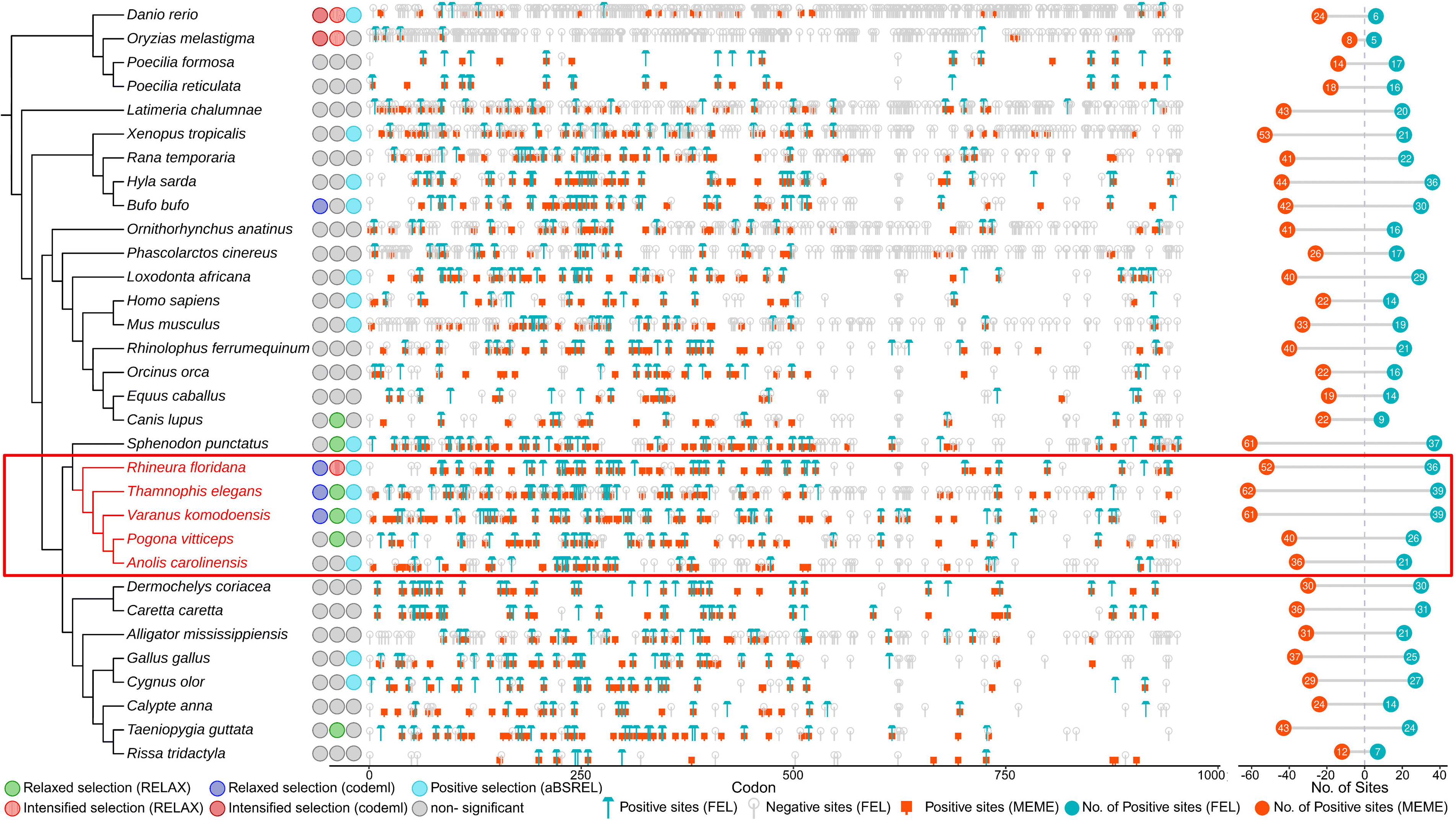
Indirect evidence for *IL34* loss: Co-evolution of receptor *CSF1R*. Selection analysis of the *CSF1R* coding sequence was conducted across 32 vertebrate species using both branch-based (RELAX, aBSREL, and CodeML) and site (FEL and MEME) models. The phylogenetic tree (left) illustrates the evolutionary relationships among species, with squamate lineages highlighted in red text and red box. Selection inferences such as relaxed/intensified and positive selection, based on RELAX, CodeML, and aBSREL analyses, are shown for each species. The central panel displays the distribution of positively and negatively selected codon sites along the gene, with codon positions on the x-axis and selection signals from focal species (foreground) against the rest of the species in the background. The right panel summarises the number of positively selected sites detected by FEL and MEME for each species, visualised using ggplot2 [124].

**Fig. 8.**
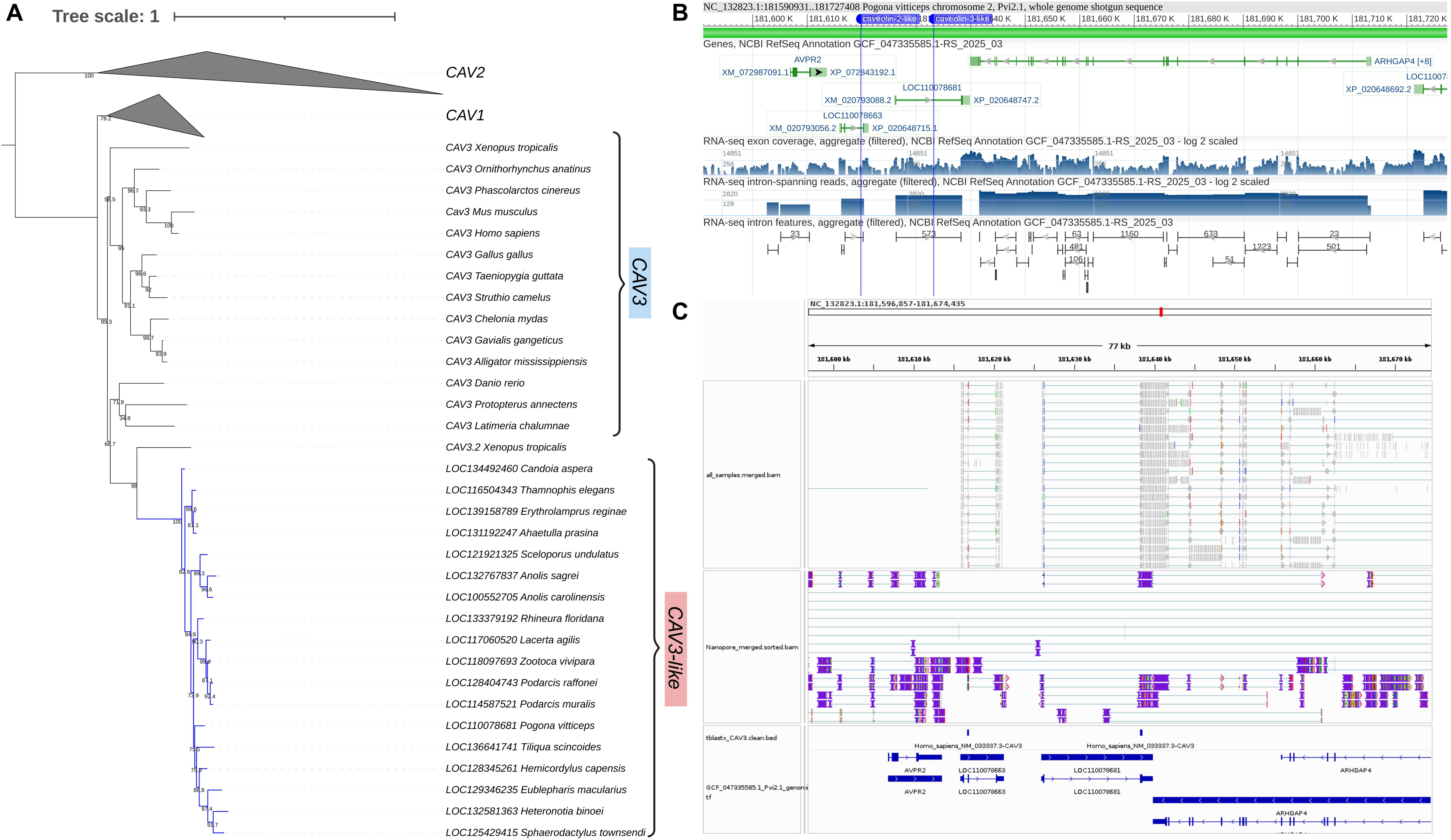
Evolutionary relationships and genomic context of caveolin (CAV) genes in Squamates. (A) Maximum-likelihood phylogeny of *CAV1*, *CAV2*, *CAV3*, and *CAV3*-like genes across representative vertebrates. *CAV1* and *CAV2* form distinct clades, while *CAV3* and *CAV3*-like (blue branch) sequences cluster separately. The phylogenetic tree made using IQ-TREE and rooted to *CAV2*. The bootstrap support values are indicated at nodes. (B) Synteny of the *CAV3*-like locus in Pogona vitticeps. (C) Transcriptomic evidence for the *CAV3*-like gene in central bearded dragon (*Pogona vitticeps*). Long-read (Nanopore) and short-read RNA-seq data confirm exon–intron structure and transcriptionally active status of *CAV3*-like (*LOC110078681*) gene in *Pogona vitticeps*.

## Discussion

Our study identifies large-scale gene losses across squamates, with 53 genes consistently unretrievable in all 261 squamate genomes screened. These genes are dispersed across macro-and microchromosomes in the chicken genome and are typically lost individually, while some are unretrievable in clusters. Based on synteny conservation, the unretrieved orthologs are classified as putatively lost due to segmental deletions (n=36) and intra/inter chromosomal rearrangements (n=17). Among the 53 unretrieved orthologs, 21 had < 25% blastn query coverage, 23 had intermediate coverage of 25–74%, and 9 had ≥75% coverage in the genomes of squamates. The mean GC content of the unretrieved orthologs ranges from 39.56 % to 64.72 %, is within a GC-neutral range and suggests that GC bias is unlikely to cause them to be unretrievable. Additionally, many of these genes exhibit less than 70% sequence identity to their human orthologues, indicating substantial evolutionary sequence divergence. Despite extensive efforts, we could not recover these genes from available genomic or transcriptomic resources, including searches in high-quality genome assemblies, analyses of raw genomic sequencing reads, and examination of RNA-seq data across diverse tissues in the central bearded dragon (*Pogona vitticeps*). The synteny and phylogenetic analyses of the 53 unretrieved orthologs revealed that 14 genes are lost in squamates with no retained paralog, 15 show ortholog loss with retained paralogs, and 24 remain unretrieved orthologs. Based on the evidence score, *IL34* is the strongest empirical case of evolutionary gene loss. Our comparative genomic analyses confirmed the complete loss of *IL34* in squamates via segmental deletion and intrachromosomal rearrangements at the *IL34* locus in some *Viperidae* snakes. We also identified squamate-specific modifications in *CSF1R*, suggesting possible compensatory changes following *IL34* loss. Similarly, *CAV3* and *KDM3A* have the highest evidence score among the 14 orthologs lost with retained paralogs. While the cross-species mapping found noisy RNA-seq support for both these genes, the phylogenetic analysis and synteny confirm the loss of the ortholog with the paralogs retained in line with the “less is more” hypothesis.

### Evolutionary loss of conserved genes in squamates: patterns and limitations

In squamates, we discovered that segmental deletions and intra- and inter-chromosomal genomic rearrangements cause the putative loss of these 53 unretrieved orthologs. In contrast with previously documented gene losses in birds, which are primarily found on microchromosomes and are found to be due to assembly errors [42], these genes are dispersed throughout different macro- and microchromosomes rather than clustered on microchromosomes in the chicken genome [38, 61]. We observed that some gene losses occurred in clusters, with two or more adjacent genes unretrievable from the same genomic region. For *IL34*, both segmental deletions and intra-chromosomal rearrangements were observed, suggesting that these loci lie in structurally unstable regions. The loss of *IL34* in squamates, along with the signatures of loss on the receptor and presence of a structural paralog [48, 59, 62], makes the case for functional compensation by distant structural homologs [63, 64]. Sequence paralogs from ancient whole-genome or segmental duplications are more widely studied than structural homologs. The example of ortholog loss with retained paralog, such as *CAV3*, *KDM3A*, *ADAP2*, *SUV39H2*, and *TBC1D14,* could be compensated for by their paralogs, such as *CAV3*-like, *KDM3B*, *ADAP1L*, *SUV39H1* and *TBC1D12.* Furthermore, the pattern of *TNIP2* presence in the vicinity of the telomeric region of chicken chromosome 4, compared to interchromosomal rearrangement in squamates (Additional file 4: Fig. S5), suggests that gene loss may be facilitated by the prevalence of the gene in highly variable and rapidly evolving regions such as telomeric and sub-telomeric regions [65, 66]. Genes such as *TNIP2* are retained as unretrieved orthologs due to the challenges of distinguishing evolutionary loss of genes from methodological limitations.

Previous studies [38, 52, 53, 61] have shown that some genes initially reported as “missing” in birds were later recovered using transcriptomic data [50], with their absence attributed to high GC content and limitations in earlier genome assemblies [49]. We incorporated GC content analysis to avoid similar errors, searched short- and long-read datasets, and examined RNA-seq data to minimise the risk of false gene loss inference in squamates. The GC content of unretrieved orthologs in squamates is less than that reported for the “missing” GC-rich genes in birds, which were later found to be “dark matter” missing in prior genome assemblies [42, 49, 67–69]. While lineage-specific GC content shifts in squamates cannot be ruled out, the presence of most unretrieved orthologs in tuatara weakens the likelihood of a widespread increase in GC content in squamates. Furthermore, we assembled gene sequences by performing BLASTn searches using raw whole-genome sequencing datasets. Phylogenetic analysis of the retrieved sequences revealed that they clustered with closely related paralogues rather than the expected orthologues. We verified genome assemblies for conserved synteny and ruled out assembly artefacts as the cause of unretrieved orthologs, suggesting that structural rearrangements, rather than assembly artefacts, may play a role in gene loss. Many of the unretrieved orthologs, including several interleukins, show less than 70% identity to their human orthologues and are primarily linked to immune functions (for gene-specific detailed references, see Additional file 1: Table S11). A plausible explanation is the involvement of these genes in host-pathogen arms races, where rapid evolution driven by changing selective pressures can lead to high sequence divergence and positive selection [70–73]. Consequently, these genes may still exist in squamates but in highly diverged forms, making them difficult to detect using standard sequence homology-based approaches. Furthermore, we could not accurately estimate deletion sizes, as the genes are absent in all the examined squamates, and the nearest outgroup with intact genes, tuatara, diverged ∼250 MYA [26]. This deep divergence limits the precise delineation of deletion boundaries, unlike in previous studies, where segmental deletions could be more clearly defined [6, 54].

The discovery of squamate-specific gene losses offers insights into their evolutionary importance, supporting the idea that gene loss can be adaptive and extending the “less-is-more” hypothesis to squamates vis-à-vis providing an alternative to phyletic gradualism [39, 65, 74–77]. Many unretrieved orthologs have multiple paralogs; in some cases, their apparent absence may result from assembly limitations. However, if the gene loss is true, its function is likely compensated by paralogous genes. Additionally, most unretrieved orthologs form conserved syntenic clusters in non-squamate vertebrates, suggesting their loss in squamates likely involved block deletions, potentially contributing to genome size reduction [37]. Future genome assemblies should aim to assess the presence or absence of these genes to understand the mechanisms and evolutionary consequences of their loss.

### Functional implications of unretrieved orthologs

Earlier studies reported the absence of *PLAAT1*, *SLC24A1*, and *IL34* in squamates and associated their loss with adaptations to low-light environments, modifications in the visual system, and the evolution of myeloid cell types, respectively [46–48]. Our methodology has extended the search genome-wide and identified these genes as unretrieved orthologs, reinforcing the reliability of our gene loss detection approach and highlighting the need for detailed functional investigations of unretrieved orthologs. A literature survey revealed that the 53 unretrieved orthologs from squamates are linked to diverse biological functions, including immunity and inflammation, cell growth, apoptosis, cancer, development, cardiovascular and metabolic processes, musculoskeletal formation, sensory function, and core cellular mechanisms like autophagy, protein degradation, and transcriptional regulation (for gene-specific detailed references, see Additional file 1: Table S11). The involvement of these genes in fundamental physiological and immune functions raises critical questions about how these pathways operate in their absence, offering a valuable opportunity to explore compensatory mechanisms and lineage-specific adaptations. The loss of these genes may play a pivotal role in evolutionary genetics by influencing genome size and complexity, affecting the rate of evolution and adaptation, offering insights into gene and pathway essentiality, and contributing to phenotypic diversity in squamates [13]. Furthermore, we found that the genes *KDM3A*, *SUV39H2*, and *RBBP7*, which are unretrieved in squamates, are associated with the Gene Ontology (GO) term “chromatin remodeling”, and are involved in DNA replication, recombination, and repair. The absence of these genes suggests a potential impact on chromatin architecture, with possible consequences for genome accessibility and stability in squamates [3]. The genes *FREM2*, *WNT2* and *NOTCH2* involved in eye development are unretrieved and may be associated with changes in the visual system in squamates [31, 46, 47].

The present study reports the loss of four genes─ *IL34*, *STAP1*, *LAPTM5* and *TNIP2-*involved in macrophage activation and polarisation. *IL34* expression is observed in tissues such as the skin, brain, kidneys, and testes and is modulated in diseases like Alzheimer’s, cancer and HBV viral infection [78–80]. The macrophages derived from *IL34* exhibit more anti-viral activity than those from *CSF1* [81–83]. Despite having a normal phenotype, *IL34*^-/-^mice exhibit impaired responses to skin antigens and viral infections, highlighting its physiological importance [84, 85]. These findings suggest that squamates may be functionally compromised owing to the loss of *IL34*, potentially rendering them more susceptible to pathogenic infections. In mice, *STAP1* deficiency impairs TCR-mediated T cell activation [86], *TNIP2* acts as a regulator of NF-κB signalling [87, 88], and *LAPTM5* inhibits HIV-1 progeny infectivity by transporting viral Env to the lysosome for degradation in macrophages [89]. However, the role of these genes in reptiles remains largely unexplored and needs functional validation. The limited availability of tools for assessing immune function in free-living reptiles presents a challenge in functionally testing this scenario. These findings offer new insights into the evolutionary reduction in the gene repertoire regulating macrophage differentiation, which may help develop novel pharmaceutical strategies for the immune modulation of monocytes and macrophages.

## Conclusions

This study systematically identified and confirmed that 53 genes are putatively lost in the squamate lineage but retained in outgroup species, suggesting lineage-specific gene loss events primarily associated with large-scale segmental deletions and chromosomal rearrangements. Among the 53 unretrieved orthologs, 14 genes are lost in squamates with no retained paralog, 15 show ortholog loss with retained paralogs, and 24 remain unretrieved orthologs. Notably, four unretrieved orthologs are involved in macrophage activation and polarisation─*IL34*, *STAP1*, *LAPTM5* and *TNIP2*, while others play key roles in immunity, development, metabolism, cellular processes, and neuronal and musculoskeletal functions. The unretrieved orthologs are GC-neutral and are members of multi-gene families with several paralogues. Our study provides direct and indirect evidence of potential gene losses in the squamate lineage. Despite comprehensive searches across genome assemblies, raw reads, and transcriptomes, none could be recovered, supporting their possible loss in squamates. Further studies are needed to generate high-quality squamate genomes to identify the causative forces behind these gene losses and their final confirmation.

## Methods

### Identification of unretrieved orthologs from squamates

We retrieved the list of all 20,595 protein-coding genes annotated in the human genome from NCBI (Additional file 1: Table S1-S2). The orthologues for each human gene across 679 annotated vertebrate genomes were obtained using the datasets command-line utility of NCBI (parameters: --ortholog all) [90]. First, as a pilot analysis to identify unretrieved orthologs specific to squamates, we focused on genes consistently identified as not annotated in NCBI RefSeq across all 30 annotated squamate genomes. As a filtering criterion to ensure that these genes were not universally absent or pseudogenised or annotation artefact, we retained only those genes that were classified as intact (annotated without the “Low quality protein” tag of NCBI) in multiple distantly related species, namely Western clawed frog (*Xenopus tropicalis*), American alligator (*Alligator mississippiensis*), chicken (*Gallus gallus*), green sea turtle (*Chelonia mydas*), and human (*Homo sapiens*) allowing to confidently infer squamate-specific unretrieved orthologs by comparing across a broad phylogenetic tree of vertebrates (Additional file 1: Table S1-S2).

### Analysis of evolutionary rearrangement and chromosomal mapping of unretrieved orthologs

To investigate the chromosomal distribution and rearranged genomic regions of unretrieved orthologs across squamates, we mapped them to chicken karyotypes. We used GENESPACE v1.3.1 [91] to understand patterns of synteny and orthology across chicken (*Gallus gallus*), American alligator (*Alligator mississippiensis*), green sea turtle (*Chelonia mydas*), central bearded dragon *(Pogona vitticeps*), brown anole (*Anolis sagrei*), and terrestrial garter snake (*Thamnophis elegans*) genomes. A comparative framework enabled the visualisation of conserved syntenic blocks and rearranged genomic regions linked to these 53 unretrieved orthologs in squamates. We used BEDTools nuc to calculate GC content across 100 bp sliding windows along the length of the chromosome. To identify sequences associated with non-B DNA-forming motifs, we used the GFA v2 program [92]. The resulting outputs were used to plot the GC content and the presence of non-B DNA motifs across chicken chromosomes. Additionally, the unretrieved orthologs loci were compared between the genome assemblies of chicken (*Gallus gallus*) and green anole (*Anolis carolinensis*) using the NCBI Comparative Genome Viewer [93].

### BLASTn search of 53 unretrieved orthologs in squamate genomes and raw read datasets

To increase the power of screening for unretrieved orthologs in squamate genomes, we performed BLASTn [94] searches (parameters: -evalue 0.05 -outfmt ‘17 SQ’, 1, 6, and 7) across 261 squamate genome assemblies, including both chromosome-level and scaffold-level assemblies, using human (*Homo sapiens*) and chicken (*Gallus gallus*) gene sequences as queries (Additional file 1: Table S5). The BLASTn output in format ‘17 SQ’ was used to generate IGV reports [95], which were then screened to assess the presence of gene sequences in the squamate assemblies. Furthermore, we screened publicly available short-and long-read DNA and RNA sequencing data from PacBio (HiFi and Revio), Oxford Nanopore, and Illumina sequencing technologies—from squamate species (Additional file 1: Table S12). We employed BLASTn (parameters: -evalue 0.05 -max_target_seqs 10000) with query sequences of unretrieved orthologs of the five outgroup species. To distinguish between orthologs and paralogs, the sequence obtained from HybPiper [96] was analysed using IQ-TREE v2.3.6 [97], along with coding DNA sequences from 39 species that produced BLASTn hits to the extracted sequence (parameters: -evalue 0.001)(Additional file: Table S12). We used the ‘ape’ package to identify the closest phylogenetic branches to the HybPiper recovered gene sequence [98]. Similar phylogenetic analysis was performed using newly available near-complete T2T genomes of the Australian central bearded dragon (*Pogona vitticeps*) [99, 100]. To identify orthologous genes across 28 species (19 squamates and 9 outgroups), protein sequences of all annotated genes in FASTA format were used as input to OrthoFinder 2 [101]. The analysis involved an all-vs-all sequence similarity search using DIAMOND, followed by clustering of genes into orthogroups, each representing a set of genes derived from a single ancestral gene in the last common ancestor. From this dataset, we focused on 53 genes of particular interest and retrieved their corresponding orthogroup IDs. The gene IDs within these orthogroups were then extracted. To confirm whether these sequences represented true orthologs, or paralogs, or gene duplicates, we conducted synteny analyses and phylogenetic inference using IQ-TREE 2 (Additional file 1: Table S13) [97].

Furthermore, to assess the paralog prevalence, we used Ensembl BioMart to obtain annotated paralogs, GC content, sequence identity, and transcript count data for human and chicken orthologs of squamate-specific unretrieved orthologs.

To explore potential transcriptional evidence for unretrieved orthologs, we mapped RNA-seq data from the central bearded dragon (*Pogona vitticeps*) of the testis, muscle, lung, liver, kidney, eye, heart, and brain tissues to the chicken genome using minimap2 v2.29-r1283 [102] (parameters: -ax splice -uf -k14) and FLAIR v2.2.0 [103] (Additional file 1: Table S12). We then screened for the presence of transcripts corresponding to unretrieved orthologs and visualised the read coverage and splice junctions using IGV-report, allowing us to assess transcriptional signals for these genes in the central bearded dragon (*Pogona vitticeps*) despite their apparent genomic absence (i.e., lack of annotation and failure to be picked up by blastn search).

### Identification of 1-to-1 orthologs, assembly verification, GC content, paralog annotation of unretrieved orthologs

We confirmed gene synteny of unretrieved orthologs and identified 1-to-1 orthologs across eight vertebrate species, namely Western clawed frog (*Xenopus tropicalis*), American alligator (*Alligator mississippiensis*), chicken (*Gallus gallus*), green sea turtle (*Chelonia mydas*), green anole (*Anolis carolinensis*), Common wall lizard (*Podarcis muralis*), Central bearded dragon (*Pogona vitticeps*), Eastern brown snake (*Pseudonaja textilis*), tiger rattlesnake (*Crotalus tigris*), tuatara (*Sphenodon punctatus*), platypus (*Ornithorhynchus anatinus*), house mouse (*Mus musculus*) and human (*Homo sapiens*) using NCBI genome viewer [22] and ENSEMBL [104]. To assess the integrity of the unretrieved orthologs loci in the chromosome/scaffold-level assembly of Indian cobra (*Naja naja*), tiger rattlesnake (*Crotalus tigris*), Central bearded dragon (*Pogona vitticeps*) and common wall lizard (*Podarcis muralis*). We used the *scan_alignment* and *alignment_plot* options in the Klumpy v1.0.10 tool [105], the required BAM file was generated by mapping long-read sequencing data from squamates species (Additional file 1: Table S12) with Minimap2 v2.17-r941 [102] (parameters: -ax map-pb or -ax map-hifi or -ax map-ont). The resulting alignment plots were examined for the presence of continuous and overlapping reads spanning the unretrieved orthologs loci.

The lack of annotations and/or sequence for the unretrieved ortholog from the genome assembly of a species could be due to its high GC content and/or high sequence divergence [42, 49, 106]. Therefore, we calculated GC content across the coding sequence of vertebrate orthologs using seqkit (parameters: fx2tab --name –gc) and sequence identity with respect to the human ortholog using EMBOSS-needle [107].

### Squamate lineage-specific evidence of segmental deletion

#### a. Dataset preparation

We developed an integrated analysis pipeline encompassing orthologue retrieval, synteny mapping, intergenic distance measurement, and gap detection to investigate segmental deletions underlying unretrieved orthologs. Firstly, four neighbouring syntenic genes (two upstream and two downstream of the focal gene) were identified with synteny analysis, and their orthologs across vertebrate genomes were retrieved from NCBI datasets (parameters: --include product-report --ortholog all). The longest isoform was selected for each ortholog based on protein length and species, where orthologs of all four syntenic genes were annotated. We calculated pairwise intergenic distances using BEDTools closest. Gene pairs on different chromosomes were flagged as potential Evolutionary Breakpoint Regions (EBRs). A comprehensive gene distance matrix with complete orthologous information was assembled for all species. Next, to detect gaps in the locus, genomic intervals spanning the outermost flanking genes were merged with BEDTools merge -d 100000000, and the corresponding sequences were retrieved using NCBI efetch. These regions were scanned for assembly gaps using the Klumpy v1.0.10 find_gaps subprogram, and the total number of gaps was recorded per species. Each species’ taxonomy classification (class and order) was obtained using NCBI Taxonomy resources (esearch and efetch). Finally, all data—including intergenic distances, gap counts, and taxonomic info—were consolidated into a unified summary table. A subset of species with complete data was used for downstream visualisation (heatmap) and phylogenetic logistic regression analysis. Based on the presence of intact synteny and absence of gaps, species were classified as having either an intact locus (non-squamates) or a potentially deleted locus (squamates).

#### b. Statistical analysis of gene distance and phylogenetic signal

##### i. Ratio-Based Comparative Analysis

To evaluate whether changes in intergenic distances between the focal gene (genes which are unretrievable in squamates) and its flanking genes are associated with segmental deletion, we first excluded species exhibiting large-scale genomic disruptions such as gaps and EBRs. We calculated six pairwise ratios representing the remaining species’ upstream, downstream, and across-gene intergenic distances. We conducted Wilcoxon rank-sum tests for each ratio with the alternative hypothesis set to “less” to compare distributions between species with intact versus deleted gene loci. The resulting *p*-values were adjusted for multiple testing using the FDR method, and significant comparisons were visualised as a heatmap using the pheatmap package in R [108].

##### ii. Phylogenetic Logistic Regression

To quantify the phylogenetic association between relative gene distance and gene loss, we performed phylogenetic logistic regression using the “phylolm” R package [109]. Squamates (0) and non-squamates (1) were used as the binary variable. The independent variable was the relative intergenic distance for the focal gene, computed as:

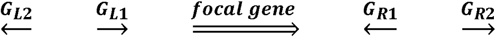

Where

*G*_*L*l_: Gene on the left flank of a lost gene on 1^St^ position

*G*_*L*2_: Gene on the left flank of a lost gene on 2^lld^ position

*G*_*R*_: Gene on the right flank of a lost gene on 1^St^position

*G*_*R*2_: Gene on the right flank of a lost gene on 2^lld^position

focal gene:unretrieved orthologs in squamates

The direction of the arrow indicates gene orientation

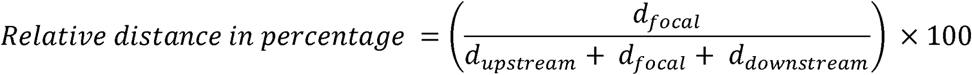

Where

*d_focal_* = distance between *G*_*L*1_ *G*_*R*1_

*d_downstream_* = distance between *G*_*L*1_ *G*_*L*2_

*d_upstream_* = distance between *G*_*R*1_ *G*_*R*2_

Two methods were used for model fitting: logistic_MPLE and logistic_IG10 (Additional file 1: Table S1).

An UpSet plot was used to visualise multiple lines of evidence supporting gene loss, including synteny-based indicators of segmental deletion or chromosomal rearrangement, genome blast coverage, validation of genome assembly integrity at the gene locus using klumpy, phylogenetic analyses for ortholog and paralog detection using whole-genome sequencing data, hybpiper & IQ-TREE, evidence of segmental deletion inferred from intergenic distances, paralog annotations from Ensembl, and orthologous sequence-based metrics such as GC content and sequence identity.

### Signatures of receptor-ligand co-evolution as indirect evidence of *IL34* loss in squamates

Structurally similar ligands can functionally compensate for the loss of each other [110]. Therefore, to explore the evolutionary impact of *IL34* loss in squamates, we investigated the *IL34*–*CSF1*–*CSF1R* signalling axis, focusing on the functional interplay between these ligands (*IL34*, *CSF1*) and their shared receptor *CSF1R*. *CSF1R* is a type III receptor tyrosine kinase critical for developing, surviving, and functioning myeloid lineage cells [111]. Previous studies have shown evolutionary co-variation between *IL34* and *CSF1R*, but not between *CSF1* and *CSF1R* [Garceau et al. [59] and reviewed in [60]]. Based on this, we hypothesised that *IL34* loss may influence the evolution and function of *CSF1R* in squamates. We leveraged ENSEMBL annotation for *CSF1R* across vertebrates and retrieved the coding DNA sequence (CDS) of the *CSF1R* with a conserved exon phase (Additional file: Table S10). Multiple sequence alignments (MSAs) of intact *CSF1R* from Squamates and vertebrate clades were generated using PRANK v.170427 in the Guidance2 suite [112, 113]. Nucleotide sequence alignments were trimmed using trimAl v1.4.1 (parameters: *-resoverlap 0.05 - seqoverlap 90*), removing sequences with >10% gaps and positions with gaps in >5% of sequences [114] and used to assess the strength of selection. Further, the alignments and phylogenetic tree obtained from the Timetree website (https://timetree.org/) were used to investigate the strength of selection [58]. We relied on the dN/dS-based method of Hyphy v2.5.48 and PAML v4.9f [115, 116]. Our selection suite consists of RELAX, BUSTED, aBSERL, MEME, and FEL frameworks of branch-site and site models of Hyphy, along with the null (M0) and branch (branch-free and branch-neutral) models of codeml [117–123]. To investigate whether the gene evolves under relaxed selection, we specified one branch as the foreground and all other branches as the background at a time in the RELAX model of the HyPhy program [68] and codeML of the PAML program [69]. A significant result of k > 1 indicates intensified selection along test branches, while k < 1 suggests relaxed selection, with significance determined at *p* < 0.05. We also implemented aBSREL and BUSTED, which are branch and gene-wide tests, respectively, to test for signatures of selection on branches of the genes. Site-specific models, such as FEL and MEME, were used to identify sites evolving under positive and negative selection, and ggplot2 was used for visualisation in the R program [124].

### Quantifying cross-species alignment conservation

To validate the absence of unretrieved orthologs in squamates, we utilised pair-wise genome alignment to quantify the alignment coverage of diverse vertebrates and compared squamates vs non-squamate orders. For this, we analysed pairwise whole-genome alignments between chicken (GRCg7b) and 27 other vertebrate species, including squamates, using precomputed LASTZ alignments from the Ensembl Comparative database (https://ftp.ensembl.org/pub/current_maf/ensembl-compara/pairwise_alignments/) [104]. Additionally, we created seven new pairwise whole genome alignments using newly available squamates genome assemblies with BUSCO > 96% (Additional file 1: Table S12). The multiple alignment format (MAF) files for each species were downloaded and processed using MafFilter v1.3.1 [125] with the OutputCoordinates filter enabled (output_src_size=yes) to extract conserved syntenic blocks between chicken and the focal species. Next, we used BEDTools intersect with the -wao option to compute overlap between each exon of 53 genes and the aligned regions for each focal species. Finally, to assess alignment for each unretrieved ortholog, we normalised the aligned region with CDS length. The resulting matrix (gene × species) of normalised aligned values was visualised as a heatmap using the pheatmaps v1.0.13 R package (Additional file 1: Table S9). Pairwise Wilcoxon rank-sum tests (alternative = “greater”) were performed to compare the aligned regions between squamates and other orders. Multiple testing correction was applied using the “BH” method, and adjusted p-values were used to assess significance (Additional file 1: Table S9). Comparisons of alignments across syntenic genes within squamates were visualised using alluvial plots and heatmaps followed by statistical evaluation using Wilcoxon rank-sum tests (alternative = “less”) to compare the left and right neighbouring genes of each focal unretrieved ortholog. Genome alignments of squamate species were assessed using both the assemblies available in Ensembl and those generated in this study.

## Data availability

The data and relevant code for this study are available on GitHub: https://github.com/CEGLAB-Buddhabhushan/Missing_genes_in_squamata.git

## Abbreviations

*EBR*: Evolutionary Breakpoint Region
*HBV*: Hepatitis B Virus
*MYA*: Million Years Ago
*TCR*: T Cell Receptor
*PacBio*: Pacific Biosciences
*ONT*: Oxford Nanopore Technologies
*FDR*: False Discovery Rate
*BH*: Benjamini–Hochberg correction
*NCBI*: National Center for Biotechnology Information
*BLASTn*: Basic Local Alignment Search Tool (nucleotide)
*MAF*: Multiple Alignment Format
*IGV*: Integrative Genomics Viewer
*RNA-seq*: RNA sequencing
*Mb*: Megabase
*Kb*: Kilobase

## Acknowledgements

We gratefully acknowledge the Vertebrate Genomes Project (VGP; https://vertebrategenomesproject.org) for generating and making available the high-quality genome assemblies that greatly facilitated this work. We thank the Council of Scientific & Industrial Research (CSIR) and the University Grant Commission (UGC) for a fellowship to BGS and SH, respectively. We used BioRender (https://biorender.com) to arrange figures and images of species.

## Author information Authors and Affiliations

Buddhabhushan Girish Salve, Sonal H, Nagarjun Vijay

Computational Evolutionary Genomics Lab, Department of Biological Sciences, IISER Bhopal, Bhauri, Madhya Pradesh, India

## Contributions

BGS and NV. conceived and designed the study; BGS, SH, and NV performed research and analysed data; and BGS, SH and NV wrote the manuscript. All authors have read and approved the final version of the manuscript and agree with the order of presentation of the authors.

## Corresponding author

Correspondence to Nagarjun Vijay

## Ethics declarations

### Ethics approval and consent to participate

Not applicable.

### Consent for publication

Not applicable.

