## Additional File 4 for "Chromosomal rearrangements and segmental deletions contribute to gene loss in squamates"

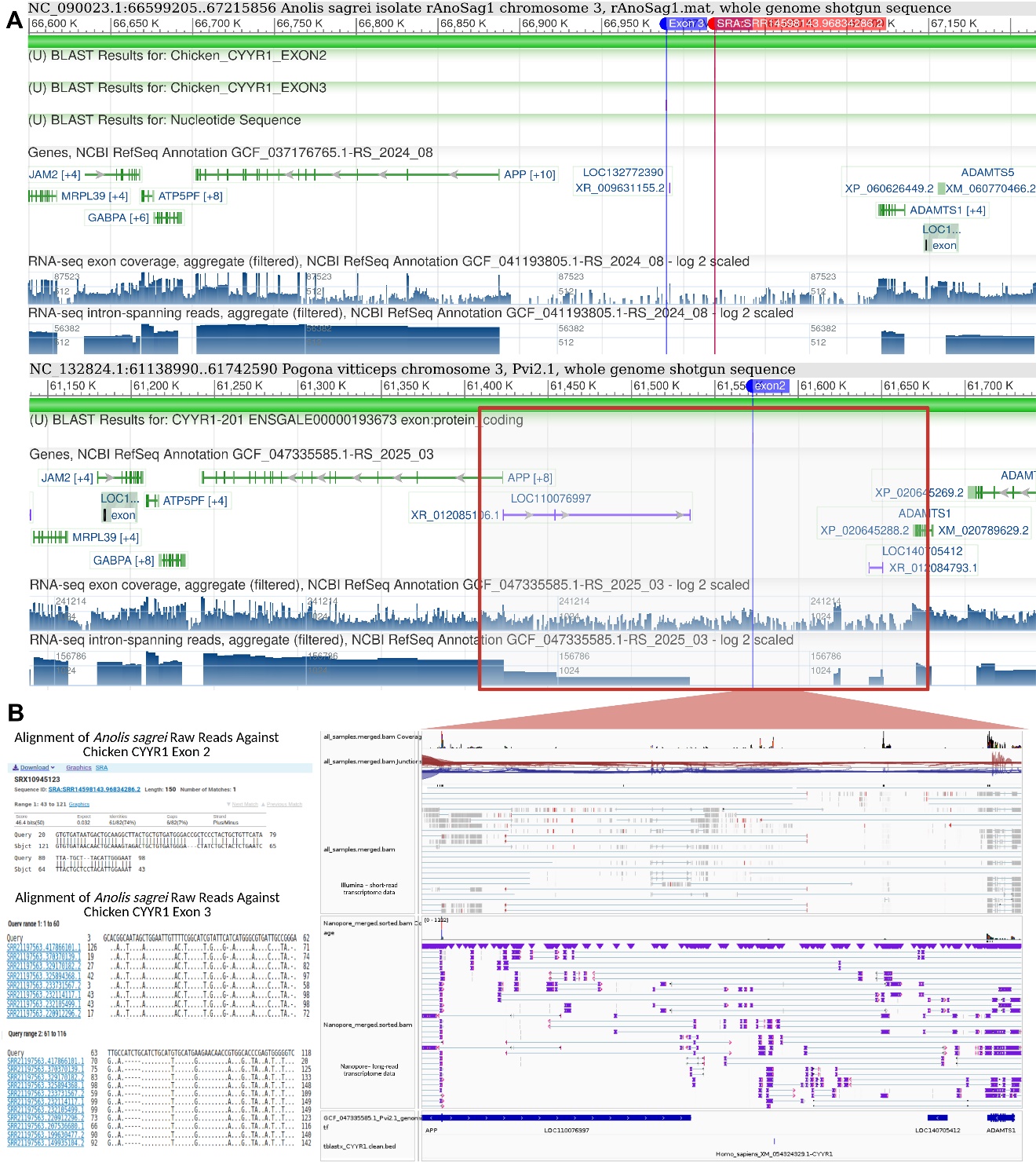

**Fig. S1** Genomic remnants of the CYYR1 gene in select squamate reptiles. **A.** NCBI Genome Viewer tracks for Anolis sagrei (top) and Pogona vitticeps (bottom). Blue tags indicate regions of significant homology to chicken CYYR1 exons 2,3, and the red tag shows the position of the raw read aligned with exon 2 within the expected syntenic locus. Although a few RNA-seq reads map to these regions (shown below, Pogona vitticeps), coverage is minimal and likely represents transcriptional noise rather than functional expression. Additional tblastx searches using the human CYYR1 nucleotide sequence fall within the expected syntenic region, as shown below the transcription track. **B.** A blastn alignment showing high nucleotide sequence conservation between chicken CYYR1 exon 2, 3 and the homologous remnant in the Anolis sagrei genome, with a 2-base deletion observed in exon 2 sequence of Anolis sagrei. Together, these results suggest that while the CYYR1 gene is largely lost in squamates, some exonic remnants persist in some species.

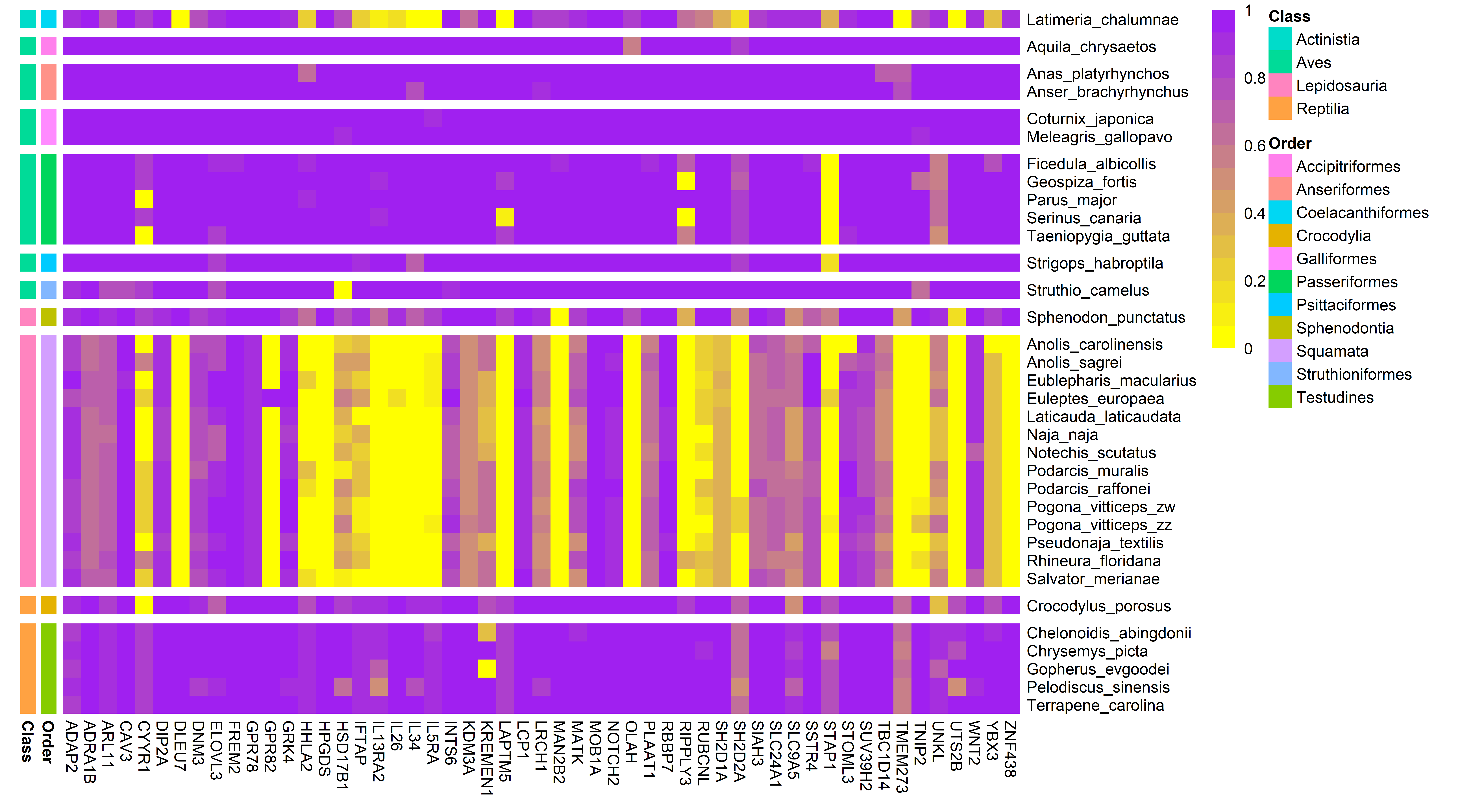

**Fig. S2.** LASTZ alignment-based evidence for unretrieved orthologs in squamates. Heatmap showing the normalised alignment of exonic regions (alignment length/CDS length) for 53 genes across 34 amniote species. Squamates exhibit consistently reduced alignment at the focal loci compared to other clades. In some cases where alignment is detected, it may originate from paralogous sequences rather than true orthologs, while unaligned regions may reflect highly divergent orthologs.

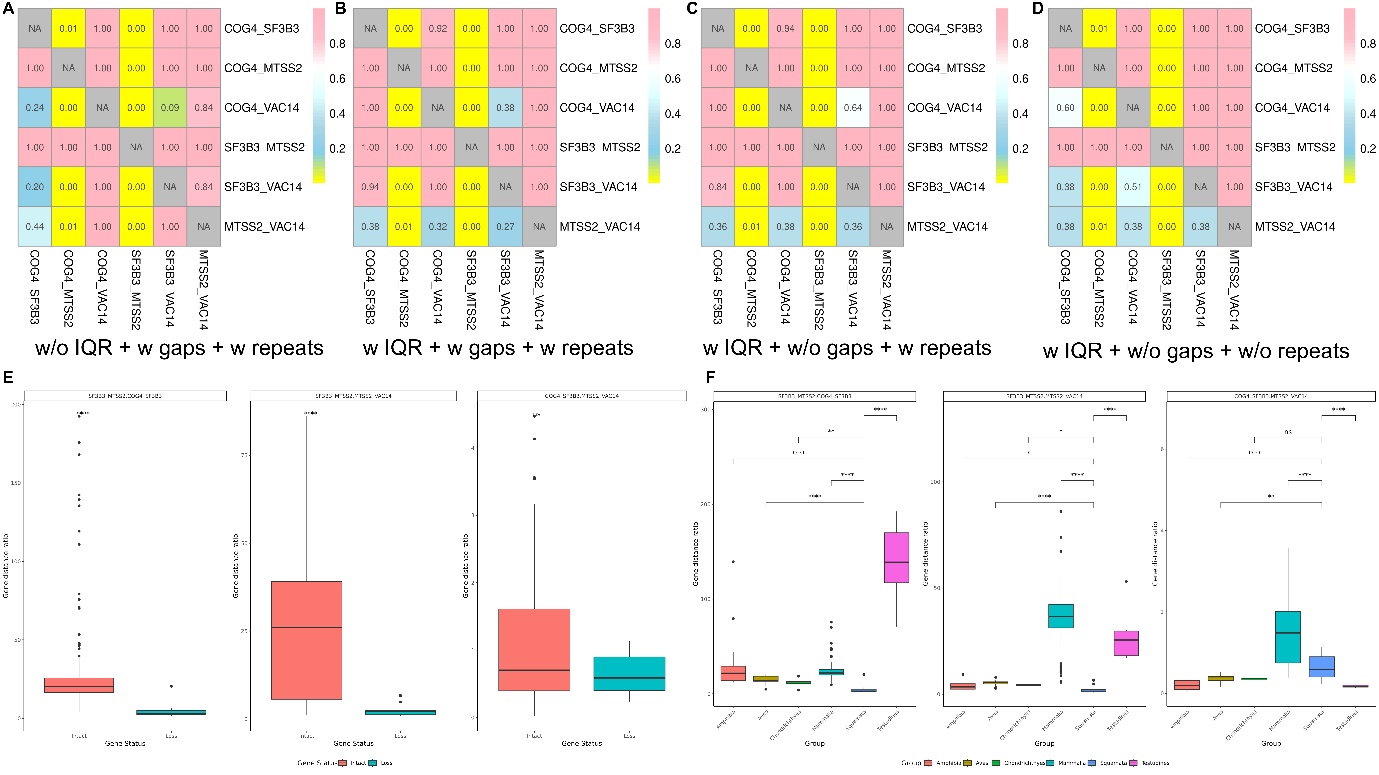

**Fig. S3** Evidence for IL34 gene loss via segmental deletion in squamates. **A-D.** The heatmap illustrates reduced intergenic distances at the IL34 locus (between SF3B3 and MTSS2) in squamates compared to non-squamates. The effect of Outliers, gaps, and repeats in the regions was evaluated to assess methodological robustness (Top panel). Across all conditions, squamates consistently show reduced gene distances at the IL34 locus. Values within the boxes indicate adjusted p-values, highlighting significant differences between squamates and non-squamates. IQR stands for interquartile range. **E.** The boxplot shows that gene distance ratios involving the IL34-spanning regions (SF3B3–MTSS2/COG4–SF3B3 and SF3B3–MTSS2/MTSS2–VAC14) are significantly lower in squamates (treated as gene loss) compared to non-squamates (intact). In contrast, no significant difference was observed for the control ratio (COG4–SF3B3/MTSS2–VAC14), which does not span the IL34 region. **F.** A similar analysis across major vertebrate groups confirms that squamates consistently exhibit reduced intergenic distances at the IL34 locus. Pairwise p-value significance from one-tailed Wilcoxon tests (alternative = "less") is shown for comparisons between squamates and non-squamates.

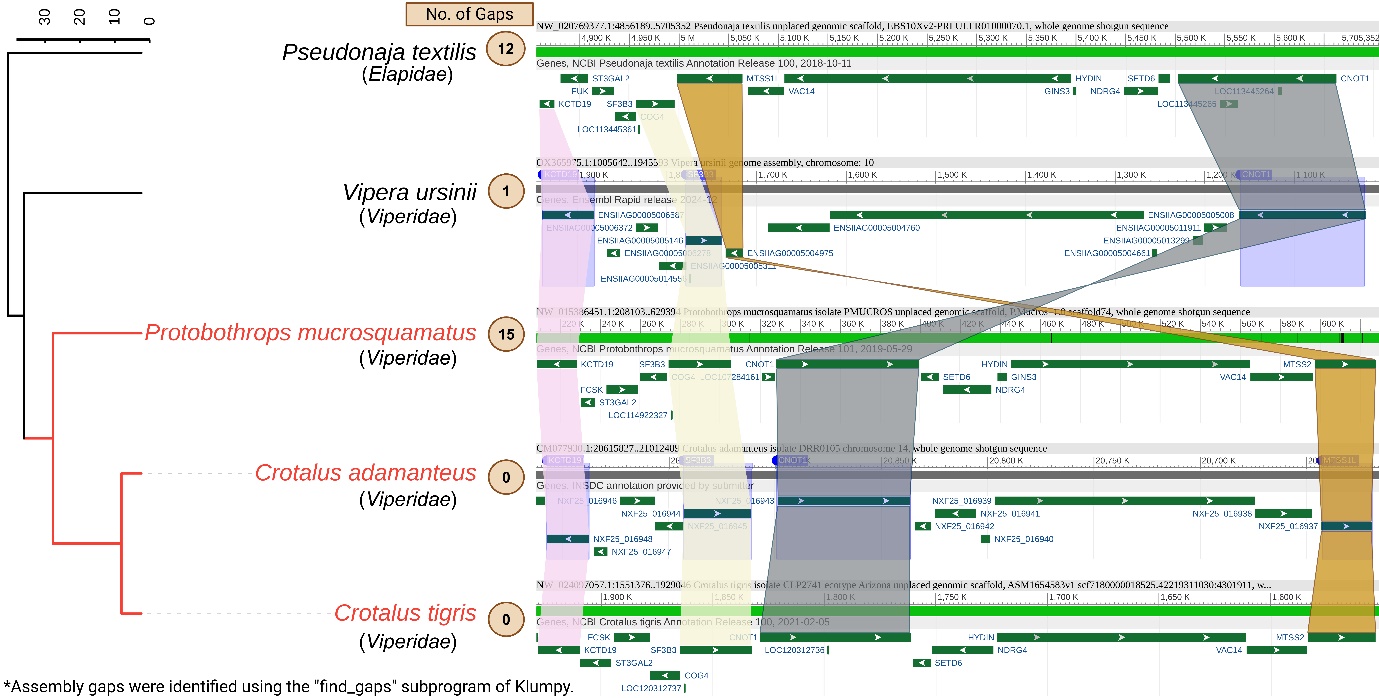

**Fig. S4** Intrachromosomal rearrangement at the IL34 gene locus in Viperidae snake species. The phylogenetic tree (left) depicts relationships among snakes (family names are shown below species names); species with intrachromosomal rearrangements are highlighted in red. Numbers within light brown circles indicate the number of gaps. Coloured ribbons connect gene orthologs, and genes are represented as green rectangular bars with arrows indicating orientation. Genomic loci were obtained using the NCBI Genome Data Viewer [1], and gaps were identified using the find_gaps subprogram of Klumpy [2]. The phylogenetic tree was obtained from the Timetree website [3] and annotated in iTOL (<https://itol.embl.de/>).

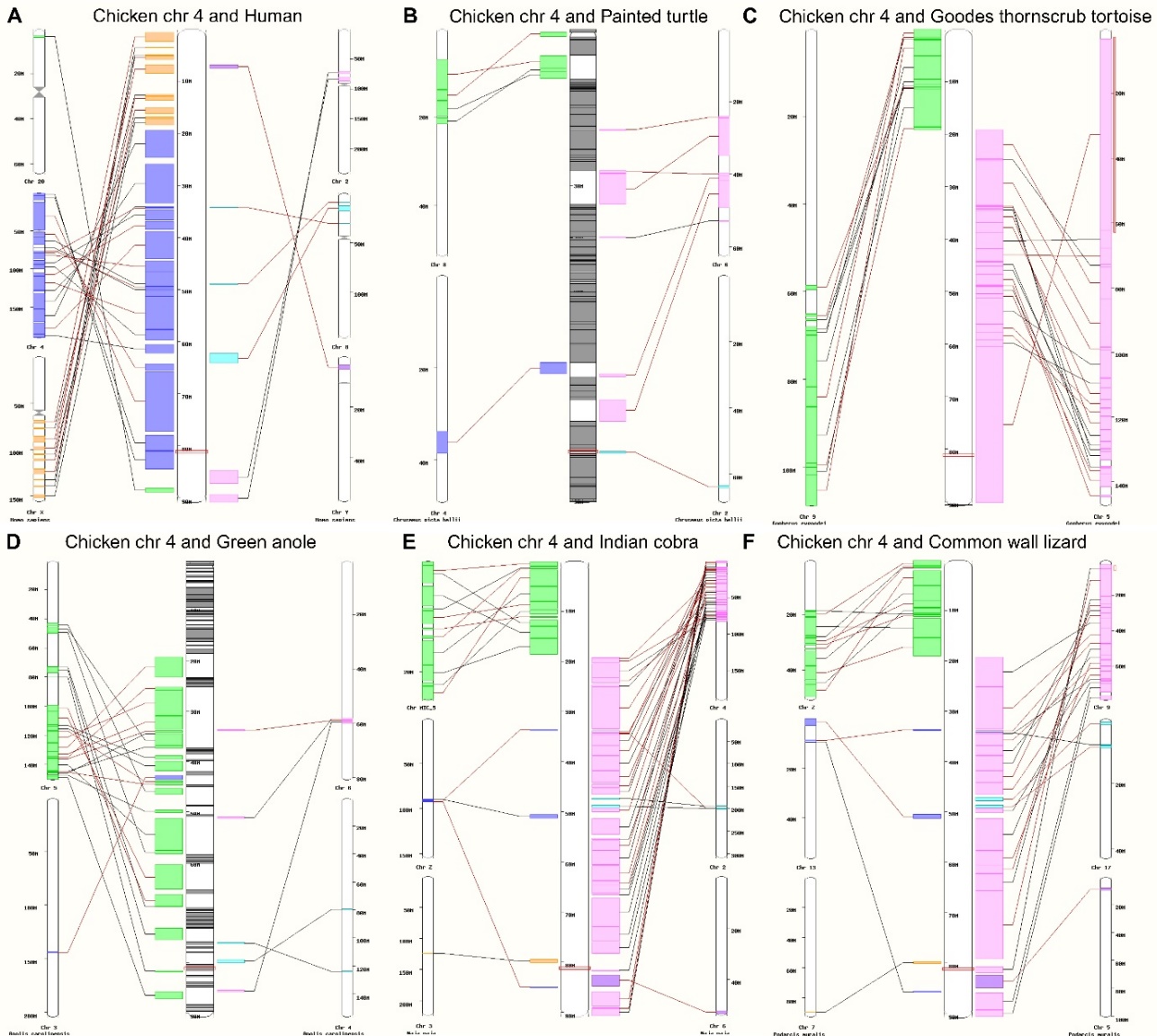

**Fig. S5** Intrachromosomal rearrangement at the TNIP2 gene locus in squamate species compared to non-squamates. The top panel shows the homologous region comparison of chicken chromosome 4 to human **A.** painted turtle (Chrysemys picta), **B**. and Goodes thornscrub tortoise (Gopherus evgoodei) **C**; in all three species, the region is projected to a single locus. The TNIP2 locus is shown in red rectangles, and homologous regions are shown in connecting lines. The bottom panel shows the red rectangle region projected to different chromosomes of squamate species, such as chromosomes 4 and 5 of green anole (**D**), chromosomes 3 and 4 of Indian cobra (Naja naja) (**E**), and chromosomes 7 and 9 of common wall lizard (Podarcis muralis) (**F**). The homologous regions are found in the telomeric and sub-telomeric regions of chromosomes. The comparison of syntenic location is done using ENSEMBL [4].
